# Multiomics analysis of narcolepsy T cells: global hypomethylation linked to T cell proliferation

**DOI:** 10.1101/2024.05.01.592019

**Authors:** Mihoko Shimada, Makoto Honda, Yoshiko Honda, Tohru Kodama, Yuki Hitomi, Katsushi Tokunaga, Taku Miyagawa

**Author notes:** Correspondence to: Taku Miyagawa, Ph.D. Department of Psychiatry and Behavioral Sciences, Tokyo Metropolitan Institute of Medical Science, 2-1-6 Kamikitazawa, Setagaya, Tokyo 156-8506, Japan. Correspondence to: Mihoko Shimada, Ph.D. Genome Medical Science Project (Toyama), National Center for Global Health and Medicine (NCGM), 1-21-1 Toyama, Shinjuku-ku, Tokyo 162-8655, Japan. These authors are the paper’s lead contacts (Mihoko Shimada, Taku Miyagawa).

## Abstract

Narcolepsy type 1 (NT1) is a chronic sleep disorder caused by a loss of orexin-producing cells in the brain and involves autoimmune mechanisms, including the presence of autoreactive T cells. We performed a genome-wide DNA methylation analysis using CD4^+^ and CD8^+^ T cells of NT1 patients. Analysis of differentially methylated regions as well as multiomics analysis with genomic and transcriptomic data obtained from the same samples indicated that cell chemotaxis pathways are implicated as a cause in the pathogenesis of NT1. Additionally, we found global hypomethylation in both the T cells of NT1 cases (CD4^+^: *P* = 1.69E-67; CD8^+^: *P* = 4.83E-12). These NT1-associated hypomethylated sites were significantly more abundant in solo-WCGW (sequences without neighboring CpGs, where W is an A or T base; *P* = 9.87E-194). Solo-WCGW tends to lose DNA methylation over the course of cell divisions, suggesting enhanced T cell proliferation in NT1.

## Introduction

Narcolepsy type 1 (NT1) is a chronic sleep disorder characterized by frequent lapses into sleep and cataplexy. Both genetic and environmental factors contribute to the development of NT1 ^1–3^. Selective loss of orexin (hypocretin)-producing cells has been observed in the lateral hypothalamus of NT1 postmortem brains, indicating that the loss of these cells is involved in the onset of NT1 ^4–6^; however, the mechanism has not yet been clarified. NT1 is strongly associated with human leukocyte antigen (*HLA*), and more than 90% of patients with NT1 carry the *HLA-DQB1*06:02* allele ^7–10^. Genome-wide association studies (GWASs) of NT1 have also identified other genetic factors, such as *TNFSF4* ^11^, *CCR1*/*CCR3* ^12^, *TCRB* ^13^, *ZNF365* ^13^*, TCRA* ^11,13,14^, *CTSH* ^11^, *P2RY11*/*DNMT1*^15^, the *IL10RB*-*IFNAR1* locus ^13^, and *CPT1B*/*CHKB* ^16^; most of these factors are immune-related. Exposure to Pandemrix, an H1N1 ASO3-adjuvanted vaccine against a pandemic influenza, was found to be an environmental factor for the development of NT1 ^17,18^, and some immunological abnormalities have been reported to trigger the development of NT1^19^. Recent studies have also reported the involvement of CD4^+^ T cells and CD8^+^ T cells in NT1. In a study using sensitive cellular screens, hypocretin-specific autoreactive CD4^+^ T cells were found in all of the NT1 patients, and hypocretin-specific autoreactive CD8^+^ T cells were found in some of the NT1 patients; in contrast, these cells were found in only a small number of the healthy controls ^20^. Another study has reported that the frequency of autoreactive CD8^+^ T cells specific for NT1-related peptides, which are presented primarily by *HLA-DQB1*06:02*, was lower in healthy individuals with the disease susceptibility allele *HLA-DQB1*06:02* than in patients with NT1 or *HLA-DQB1*06:02*-negative healthy individuals ^21^. In addition, it was reported that a higher percentage of CD4^+^ and CD8^+^ T cells release cytokines in response to hypocretin in children with NT1^22^, and that variations in the memory subsets of CD4^+^ and CD8^+^ T cells are higher in patients with NT1 ^23^. These results suggest that an autoimmune basis involving CD4^+^ and CD8^+^ T cells is associated with the pathogenesis of NT1.

DNA methylation is one of the major epigenetic mechanisms. The DNA methylation status affects gene expression, and it has been investigated in many diseases, because it is a relatively stable modification (although it can be changed by some factors) and is more readily measurable than gene expression, which is susceptible to various factors and prone to change ^24,25^. Causal mutations in DNA methyltransferase 1 (*DNMT1*) have been found in families with atypical narcolepsy ^26^, and it has been reported that patients with *DNMT1* mutations show a genome-wide methylation abnormality ^27^. In addition, a GWAS of NT1 has reported a significant association between single-nucleotide polymorphisms (SNPs) near *DNMT1* and NT1 ^15^. We previously performed an epigenome-wide association study (EWAS) of DNA methylation using peripheral blood samples, and we found that methylation around *CCR3* (C-C motif chemokine receptor 3) and genes related to hormone secretion and monocarboxylic acid metabolism pathways were associated with NT1; in addition, we also found that NT1-related methylation sites were significantly located outside of CpG islands, 95% of which were hypomethylated in NT1 ^28^. The array we used in that study was the Infinium HumanMethylation27 BeadChip (27K array; Illumina, San Diego, CA, USA), which could only measure methylation in about 27,000 locations. Although DNA methylation level is reportedly correlated with methylation status in neighboring sites, the correlation is not so strong as the linkage disequilibrium of SNPs ^29^, and the analysis of 27,000 sites was considered to be insufficient for understanding the entire genome. Furthermore, the DNA methylation patterns are known to vary widely among cells and tissues ^30^, and our previous study using only peripheral blood cells might have been insufficient for detecting subtle changes in methylation that occur in individual immune cells.

In the present study, we performed EWASs of DNA methylation using CD4^+^ and CD8^+^ T cells isolated from whole blood samples to detect NT1-related differentially methylated sites, as these two types of immune cells have been suggested to play an important role in the pathogenesis of NT1. We also performed RNA sequencing (RNA-seq) using whole blood from the same subjects for methylation analysis to examine NT1-associated gene expression, at least that at the time of blood sampling. Genome-wide genotyping was also carried out, and we performed multi-omics analysis using genome, methylome, and transcriptome data collected from the same subjects. Although gene expression is the most phenotypic of these omics data, we focused our analysis on methylation with consideration of the potential of DNA methylation as a form of epigenetic memory ^31^. Although it is difficult to capture the gene expression changes that trigger the development of NT1 in real-time since there is an inevitable time lag between disease onset and diagnosis (including sleep studies and blood sampling), DNA methylation may be a particularly useful tool for studying disease onset as the effects of such abnormal immune events may be retained as epigenetic memory.

## Results

### DNA methylation analysis and characteristics of the samples based on DNA methylation

We examined the DNA methylation profiles of CD4^+^ and CD8^+^ T cells using the EPIC methylation array. Our research design is shown in Fig. 1. After probe filtering and data normalization, 746,913 probes were selected for the following analysis (Fig. S1). We performed principal component analysis (PCA) analysis using normalized β values for all methylation sites, and confirmed that the CD4^+^ T cell-derived and CD8^+^ T cell-derived samples fell into widely different clusters, and that there was no significant bias in the distribution of the cases and controls within each cluster (Fig. S2A, B). We next examined whether the methylation rate of the master transcription factor binding site of T cells explained most of the variations in the PCA, as has been reported in previous studies ^32^. Variation in PC1 was found to be well-explained by the methylation rate of the T-bet binding sites, and variation in PC2 was found to be well-explained by the methylation rate of the GATA3 binding sites (Fig. S2C, D). These results indicated that the variability in the present sample mainly reflected differences in the T cell subtypes. Although the PCA results showed no relationship between T cell subtype variation and case or control bias, we further examined in more detail the presence of T cell subtypes within each of the CD4^+^ and CD8^+^ T cell samples by estimating the composition of these subtypes in the discovery set. Estimation using the MeDeCom method revealed the lowest cross-validation error when assuming six subtypes for CD4^+^ T cells and four subtypes for CD8^+^ T cells (Fig. S3). We then compared the proportion of each estimated subtype between the NT1 cases and controls, and found that only the proportion of latent methylation component (LMC) 4 in the CD4^+^ T cell analysis was significantly smaller in the NT1 cases than in the controls (*P* = 0.0032; Fig. S4A, B). Furthermore, we verified whether the samples analyzed as CD4^+^ and CD8^+^ T cells were correctly estimated to be the respective T cells using a reference-based estimation method. The results showed that in most of the samples, the estimated T cell types (CD4^+^ T cells or CD8^+^ T cells) were largely consistent with the sorted T cell types, but in three control samples of CD8^+^ T cells, we found that CD8^+^ T cells were not estimated to be the major component since there was a higher percentage of other leukocytes (Fig. S4C). These three control samples were excluded from the subsequent association analyses, because they could not be properly evaluated for CD8^+^ T cell methylation. Similarly, estimation of the T cell proportions was also performed for the replication samples. In five NT1 samples and four control samples, we found that CD8^+^ T cells were not correctly estimated to be the major component, and these samples were thus excluded from the subsequent association analyses (Fig. S4C).

**Figure 1.**
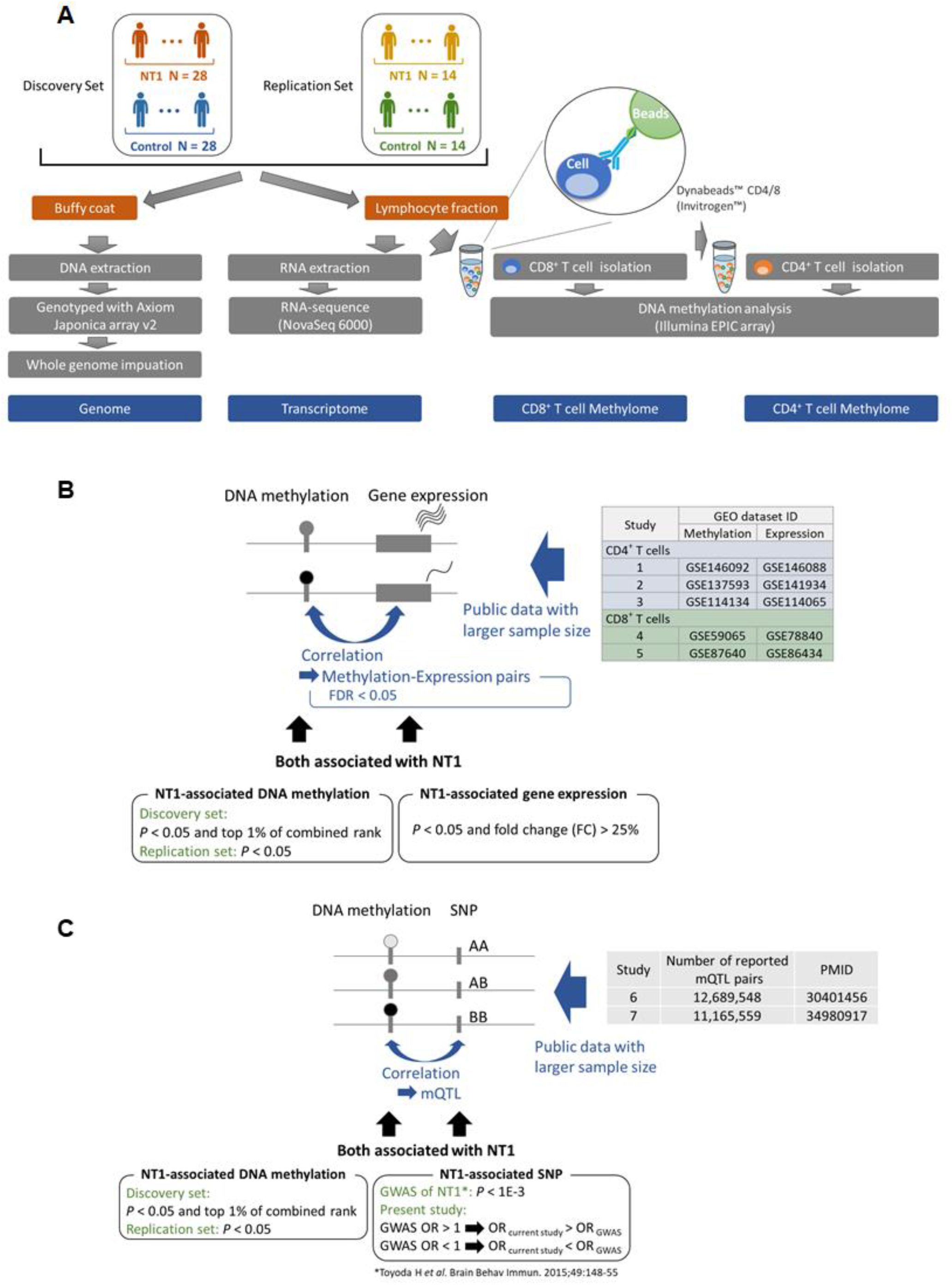
Overview of the study. **A.** Study design and graphical abstract for multiomics analysis. The discovery set consists of 28 cases of NT1 and 28 controls, while the replication set consists of 14 cases of NT1 and 14 controls. From each sample, the buffy coat and leukocyte fractions were collected. DNA was extracted from the buffy coat and genotyped using an array. From one of the two divided leukocyte fractions, RNA was first extracted, then RNA sequencing was performed to obtain transcriptome data. Using the remaining leukocyte fraction, CD8^+^ T cells were isolated using beads, followed by the isolation of CD4^+^ T cells. DNA was extracted from each of these isolated T cell types, and the DNA methylation rate was measured using an array. **B.** Methodology of the integrative analysis of methylome and transcriptome data. Initially, we identified pairs of methylation rate and gene expression that correlated with each other (Methylation-Expression pairs) using data from previous studies with larger sample sizes. Among the identified pairs, we then explored pairs where both the methylation rate and gene expression were associated with NT1 in our EWAS and RNA-seq. **C.** Methodology of the integrative analysis of methylome and genome data. From among the mQTLs detected in previous studies with larger sample sizes, we searched for mQTLs that had both a methylation rate and genotype that were associated with NT1 in our EWAS and GWAS.

We also estimated the methylation age in the discovery set, and confirmed a strong correlation between the DNA methylation age and chronological age (r = 0.913; Fig. S5A, B, C). The biological age was not increased in the NT1 cases when compared to the controls (Fig. S5C). The DNA methylation age and chronological age were also strongly correlated in the replication samples (r = 0.949; Fig. S5D).

### Differentially methylated positions and regions of NT1

We devised two strategies to identify methylation associated with NT1 to minimize false positives. One was to identify differentially methylated regions (DMRs), which are not just single-point associations like differentially methylated positions (DMPs), but regions that have a chain of related methylation sites. The other was to investigate the DMPs that have been indicated to be associated with NT1 by other omics data.

First, we examined DMPs to explore the overall trends of the associations. We found DMPs that were significantly associated with NT1 after Bonferroni correction and were replicated with *P* < 0.05 in three loci in the CD4^+^ T cell analysis; they were all localized to methylation sites within the *HLA* region (Fig. 2A), and no corresponding significant DMPs were detected in CD8^+^ T cell analysis (Fig. 2B). In our analysis, probes with poor performance were excluded; however, the *HLA* region, being highly polymorphic and challenging to analyze due to variations in probe affinity caused by the polymorphisms, was analyzed separately, and the findings have been reported in a dedicated manner elsewhere ^33^. Despite the lack of substantial differences in the proportions of CD4^+^ T cell and CD8^+^ T cell subtypes between the case and control groups, inflation of the observed *P*-values was evident in the quantile-quantile (Q-Q) plots for the analyses involving both CD4^+^ and CD8^+^ T cells. When we categorized differentially methylated sites into those that were hypermethylated or hypomethylated in NT1, it was discernible that the inflation was more pronounced at sites of hypomethylation (Fig. 2C, D).

**Figure 2.**
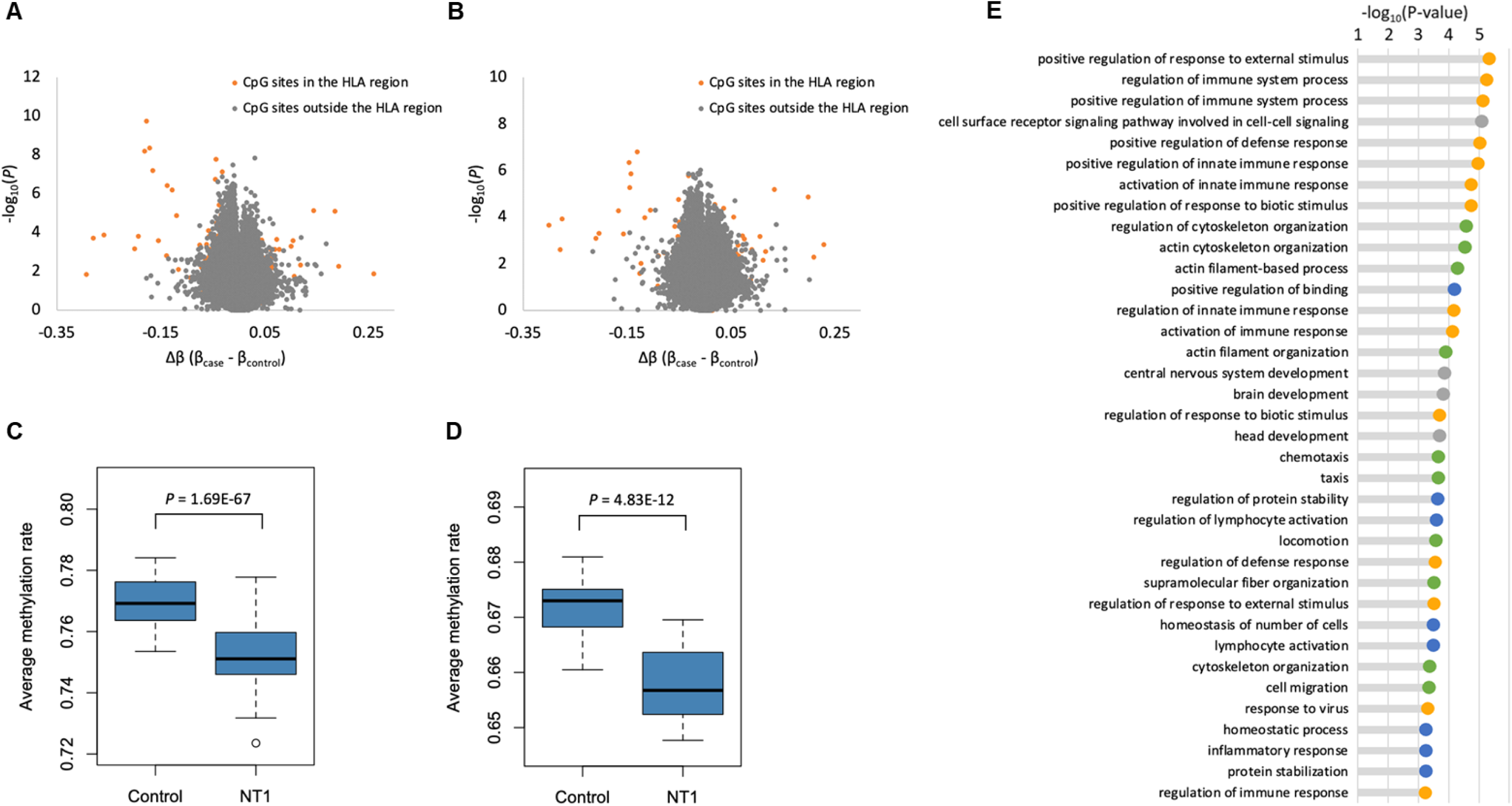
Results of the association analysis in CD4^+^ and CD8^+^ T cells. **A.** Volcano plot of the DMP analysis in the CD4^+^ T cells. **B.** Volcano plot of DMP analysis in the CD8^+^ T cells. **C.** Q-Q plot of the DMP analysis in the CD4^+^ T cells. We plotted the methylation sites separately, categorizing them those that are hypomethylated and those that are hypermethylated in NT1. **D.** Q-Q plot of the DMP analysis in the CD8^+^ T cells. Similar to the CD4^+^ ll analysis, we plotted the results of the methylation sites separately as hypomethylated or hypermethylated in NT1. **E.** Results of the pathway analysis g genes suggested to be associated with NT1 in CD4^+^ T cells. For easier interpretation, pathways were clustered into four groups using the k-means hod. The groups were color-coded as follows: yellow, pathways related to the immune response; green, pathways related to cell migration; gray, neural ways; and blue, others.

Next, we focused our investigation on DMRs to identify plausible methylation sites associated with NT1. We identified 15 replicated DMRs in the CD4^+^ T cell analysis, and five replicated DMRs in the CD8^+^ T cell analysis (Table 1). Of the DMRs detected in the CD8^+^ T cell analysis, three were also identified in CD4^+^ T cells. One of these three DMRs was located in a region extending from upstream to the first exon of the C-C motif chemokine ligand 5 (*CCL5*) gene on chromosome 17 (Table 1). Another was located in an area near the cytoplasmic linker associated protein 2 (*CLASP2*) gene, which included a methylation site that was found to be associated with C-C motif chemokine receptor 4 (*CCR4*) gene expression in CD4^+^ T cells (Table 1).

**Table 1.**
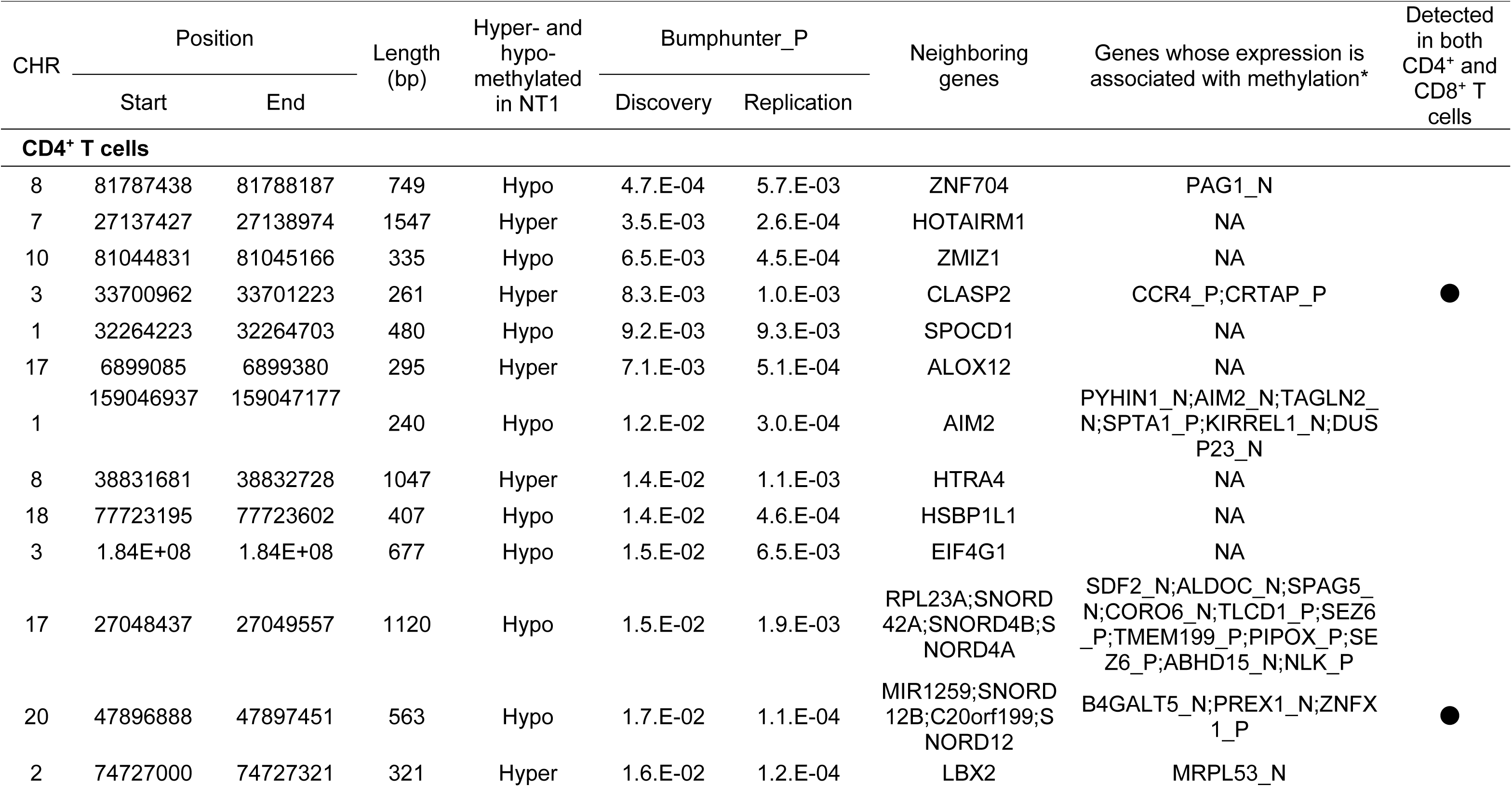
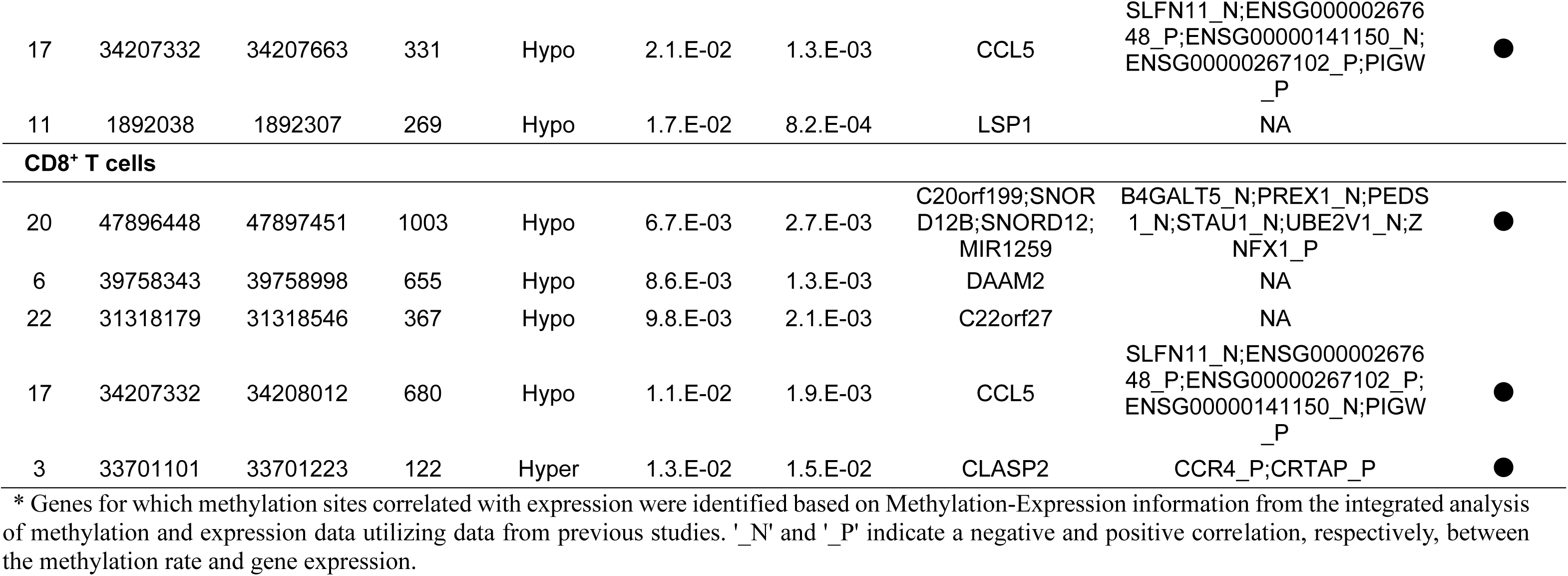
DMRs in NT1 in the CD4^+^ and CD8^+^ T cell analyses.

### Integrative analysis of methylation and gene expression

To reduce type 1 errors in the detection of DMPs associated with NT1 from genome-wide data, we examined DMPs that have been indicated to be associated with NT1 by gene expression (Fig. 1B). As it is well-known that the expression levels of numerous genes are related to the DNA methylation levels of nearby methylation sites, we first searched for methylation status and gene expression pairs that correlated with each other (Methylation-Expression pairs) from previous studies with larger samples sizes that examined both DNA methylation and gene expression in CD4^+^ and CD8^+^ T cells throughout the entire genome (Table S1) ^34–38^. The results of the EWAS for DMPs indicated that the majority of NT1-related methylation sites are common to both CD4^+^ and CD8^+^ T cells, although the strength of the association with NT1 was stronger in CD4^+^ T cells (Fig. 2); consequently, our analysis mainly focused on CD4^+^ T cells. We analyzed the results for CD4^+^ T cells from the three previous studies (Study 1, 2, and 3) listed in Table S1. In total, 3,191 and 78,881 Methylation-Expression pairs with a false discovery rate (FDR) < 0.05 were detected in Study 1 and 2, respectively. In Study 3, results for both naive CD4^+^ T cells and CD4^+^ T cells after stimulation were available, so we analyzed both, and detected 4,272 and 26,725 Methylation-Expression pairs, respectively (Table S1). These Methylation-Expression pairs were combined for a total of 110,713 Methylation-Expression pairs. Utilizing these pairs, we searched for combinations where both the methylation status and gene expression were associated with NT1, and the direction of this relation aligned with the direction of the relation in the Methylation-Expression pairs detected from the previous studies. We set the following criteria for each analysis: NT1-related methylation with *P* < 0.05 and a combined rank within the top 1% in the NT1 EWAS for DMP; and NT1-related gene expression with *P* < 0.05 and fold change (FC) > 25% in the RNA-seq analysis. As a result, four Methylation-Expression pairs were detected (Table S2). Among these, two pairs were related to natural killer (NK) cell granule protein 7 (*NKG7*), and the two other pairs were related to programmed cell death 1 (*PDCD1*) and junctional adhesion molecule 2 (*JAM2*). All of these genes were genes that showed a negative correlation between their expression levels and the methylation levels of NT1-associated methylation sites, and in the NT1 cases, the expression levels of all of these genes increased and the methylation rate at the methylation sites decreased.

Although we also conducted an analysis for CD8^+^ T cells, there were no previous studies with data were available on the detection of methylation using whole-genome bisulfite sequencing or the EPIC array and the measurement of gene expression. Therefore, we conducted the analysis of CD8^+^ T cells using the results of two previous studies (Study 4 and 5) that utilized the Infinium Human Methylation 450K BeadChip (450K array; Illumina; Table S1), which preceded the EPIC array (850K array), making it difficult to compare the CD8^+^ T cell analysis results with the CD4^+^ T cell analysis results. When the analysis was conducted under conditions similar to those for the CD4^+^ T cell analysis, no Methylation-Expression pair that met the criteria was detected in the CD8^+^ T cell analysis. We then relaxed the conditions for detecting correlated Methylation-Expression pairs from FDR < 0.05 to a nominal significance level of *P* < 0.05, and conducted the analysis. As a result, we identified three Methylation-Expression pairs, one of which was related to *NKG7,* which was also detected in the analysis of CD4^+^ T cells. Similar to the CD4^+^ T cell analysis, its expression level was increased in the CD8^+^ T cells in NT1 (Table S3).

### Integrative analysis of methylation and genotype data

To integrate the genotype and methylation data, we used information of the methylation quantitative trait locus (mQTL; Fig. 1C). Although tissue- and cell-specific mQTLs were present, in the context of QTL analysis, it has been demonstrated that larger sample sizes enhance the reliability of the results and enable the detection of a greater number of QTLs ^39^. Furthermore, prior research has reported that 84% and 80% of the effects of mQTLs detected in CD4^+^ and CD8^+^ T cells, respectively, exhibit directions concordant with those detected in whole blood. Consequently, we employed data pertaining to mQTLs identified in studies focusing on whole blood with sample sizes exceeding 1,000, as detailed in Table S4, for our integrative analysis. As shown in Table S4, Study 6 and 7 reported 12,689,548 and 11,165,559 mQTLs, respectively. Among these mQTLs, we searched for mQTLs related to NT1 that met the following conditions for both the involved SNP and methylation: for the SNP, a *P* < 1E-3 in the GWAS of NT1 ^12^ and an odds ratio (OR) that was in the same direction and greater than the OR in the GWAS in the samples of the present study (discovery and replication samples, NT1: N = 42 and control: N = 42); and for the methylation, a combined rank within the top 1% in the discovery samples that was replicated. As a result, in CD4^+^ T cells, four mQTLs from Study 6 and two mQTLs from Study 7 were detected, with no mQTLs commonly detected in both studies (Table S5). The four mQTLs detected in Study 6 were all pairs consisting of two nearby SNPs and two methylation sites, which were found to negatively correlate with the expression of S100 calcium binding protein A4 (*S100A4*; Table S5).

When a similar analysis was conducted for CD8^+^ T cells, no mQTL showing relationships between NT1 and both methylation and SNP was found. When the criteria for SNPs related to NT1 in the GWAS were relaxed from *P* < 1E-3 to *P* < 0.01, two mQTLs were detected (Table S6). However, in CD8^+^ T cells, the correlation of SNPs and methylation pairs was weaker than that in the CD4^+^ T cells.

### Pathway analysis of genes affected by methylation sites associated with NT1

Through the detection of DMRs and the integrative analysis of gene expression and genotype data, various associated methylation regions or sites were identified, especially in the analysis of CD4^+^ T cells. Consequently, we conducted a pathway analysis of genes associated with the methylation rates at these methylation sites by using the Methylation-Expression information obtained from the CD4^+^ T cell integrative analysis. The results revealed the involvement of pathways related to immune responses, chemotaxis, and cell migration (Fig. 2E and Table S7).

### Characterization of NT1-related hypomethylated sites

In the process of conducting the association analyses for the identification of DMPs, a significant inflation in *P*-values was observed across the analyses of both cell types, with this phenomenon being more pronounced in CD4^+^ T cells. Additionally, it was found that this inflation predominantly occurred at methylation sites exhibiting hypomethylation in NT1. Therefore, we explored the characteristics of these hypomethylated sites in NT1 to determine why such a pronounced trend of hypomethylation was observed in NT1.

In the examination of the prevalence of hypomethylation, the results indicated that for the discovery set, across all thresholds ranging from a nominally significant *P* = 0.05 to 1E-5, more than 97% of the replicated NT1-associated methylation sites exhibited hypomethylation in the CD4^+^ T cells (Table 2). In contrast, 61% of all analyzed methylation sites were hypomethylated in NT1. When comparing this ratio to the proportion of NT1-associated methylation sites that were hypomethylated in NT1, the results showed that the latter proportion was significantly larger (discovery set: probes with *P* < 0.01; *P* = 1.69E-67; Table 2). A similar trend was observed in CD8^+^ T cells, in which over 80% of the NT1-associated methylation sites were hypomethylated in NT1 (Table 3). We then examined the regions in which NT1-associated hypomethylated sites were common with a *P* < 0.01 in the discovery set and were replicated with a *P* < 0.05. In both T cell analyses, we found that NT1-related hypomethylated sites were significantly less frequent within CpG islands (CD4^+^ T cells: *P* = 3.03E-115 and CD8^+^ T cells: *P* = 9.30E-34), and that they became more frequent with increasing distance from the CpG islands in the shores and shelves (Fig. 3A and Table S8). Furthermore, NT1-associated hypomethylated sites were particularly scarce in exon and promoter regions (exons, CD4^+^ T cells: *P* = 6.70E-26 and CD8^+^ T cells: *P* = 7.98E-13; Promoter_inactive, CD4^+^ T cells: *P* = 2.99E-22 and CD8^+^ T cells: *P* = 1.61E-09; Fig. #B and Table S9). When compared to the related regions from ChIP-seq in the T cells, NT1-related hypomethylated sites were significantly less frequent in regions marked by histone modifications associated with transcriptional activation, such as H3K4me3 and H3K27ac (Fig. #C and Table S10). Interestingly, despite the low overlap between NT-associated hypomethylated sites in the CD4^+^ T cell analysis (N = 3,259) and those in the CD8^+^ T cell analysis (N = 897), with only 38 sites in common, both shared the characteristic that they were less common in regions likely to influence gene expression, such as CpG islands, promoter regions, and open chromatin.

**Figure 3.**
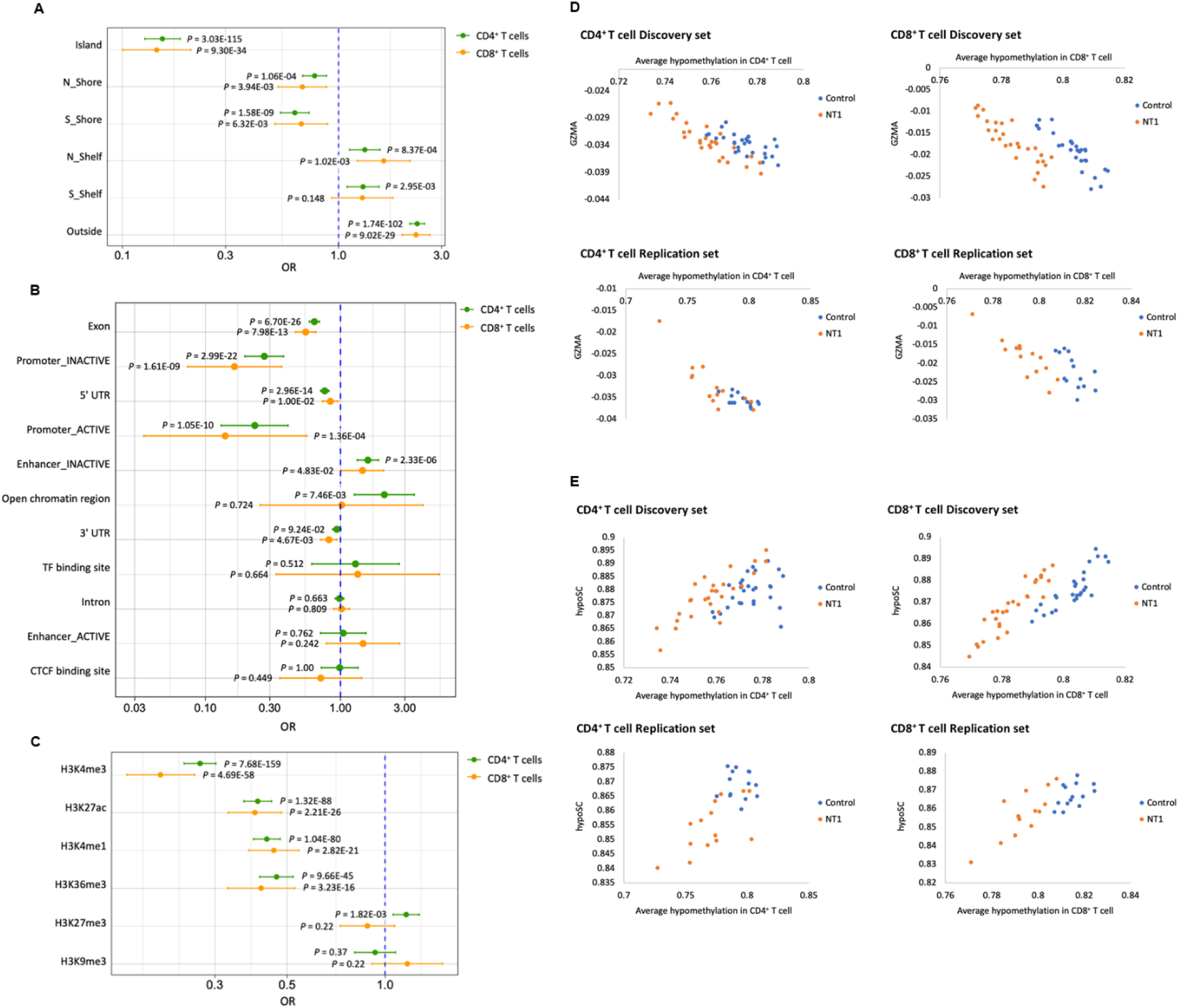
Regions with a high frequency of NT1-related hypomethylation sites and indices that showed a correlation with the degree of hypomethylation. A. The relationship between NT1-related hypomethylation sites and CpG islands. B. The relationship between NT1-related hypomethylation sites and gene regions. Regarding promoters and enhancers, categories other than “active” and “inactive” have been omitted from the figure, and the detailed results are described in Table S8. C. The relationship between NT1-related hypomethylation sites and regions marked by histone modifications. D. Correlation between the estimated values of GZMA and the degree of hypomethylation (adjusted for age). E. Correlation between hypoSC and the degree of hypomethylation (adjusted for age).

**Table 2.**
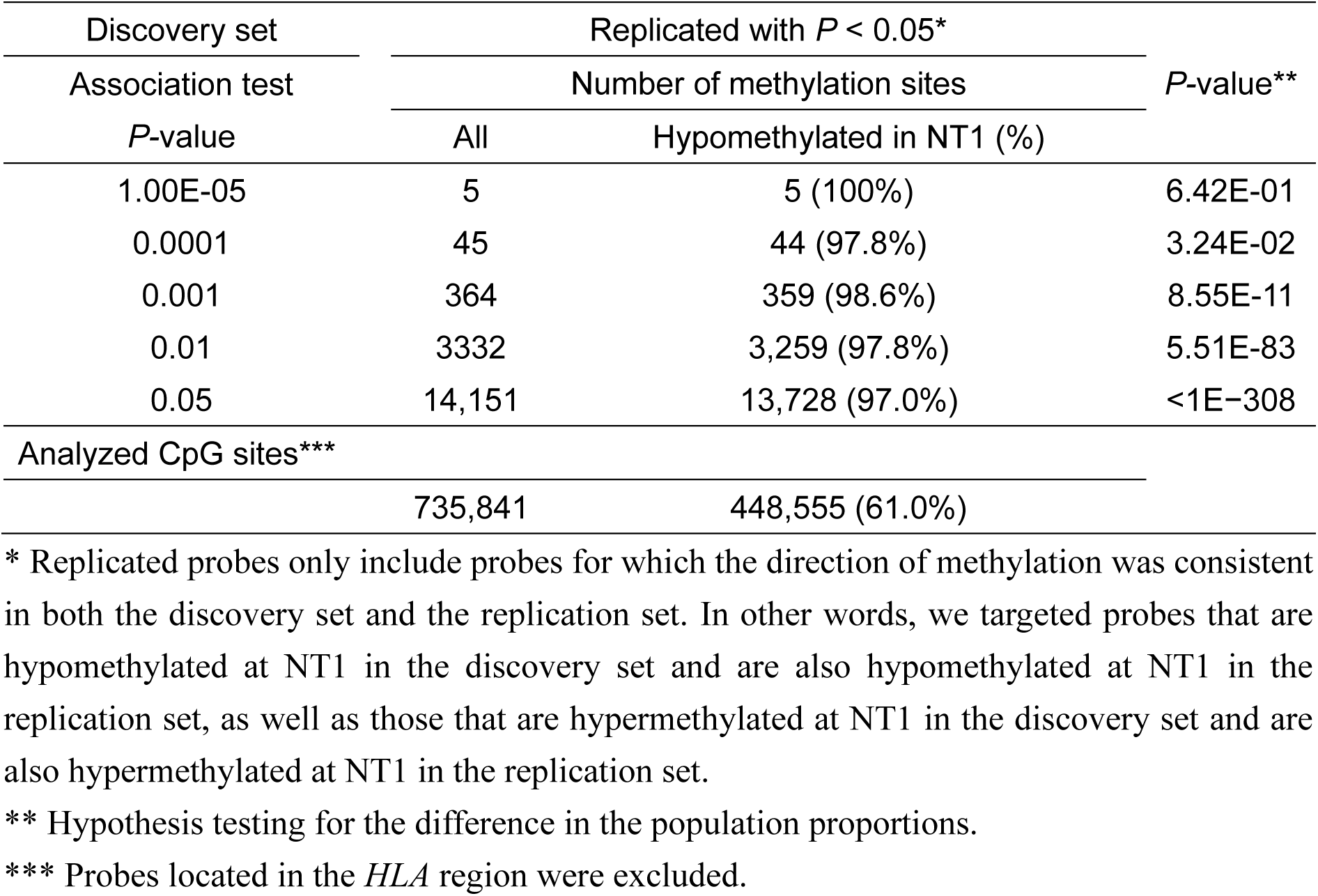
The proportion of hypomethylated sites among the NT1-related methylation sites in CD4^+^ T cells.

| Discovery set | Replicated with $P < 0.05^*$ | | $P$-value** |
| --- | --- | --- | --- |
| Association test | Number of methylation sites |  |  |
| $P$-value | All | Hypomethylated in NT1 (%) | |
| 1.00E-05 | 5 | 5 (100%) | 6.42E-01 |
| 0.0001 | 45 | 44 (97.8%) | 3.24E-02 |
| 0.001 | 364 | 359 (98.6%) | 8.55E-11 |
| 0.01 | 3332 | 3,259 (97.8%) | 5.51E-83 |
| 0.05 | 14,151 | 13,728 (97.0%) | <1E-308 |
| Analyzed CpG sites*** |  |  |  |
|  | 735,841 | 448,555 (61.0%) |  |

\* Replicated probes only include probes for which the direction of methylation was consistent in both the discovery set and the replication set. In other words, we targeted probes that are hypomethylated at NT1 in the discovery set and are also hypomethylated at NT1 in the replication set, as well as those that are hypermethylated at NT1 in the discovery set and are also hypermethylated at NT1 in the replication set.
\*\* Hypothesis testing for the difference in the population proportions.
\*\*\* Probes located in the *HLA* region were excluded.

**Table 3.** The proportion of hypomethylated sites among the NT1-related methylation sites in CD8^+^ T cells.

| Discovery set | Replicated with $P < 0.05^*$ | | $P$-value** |
| --- | --- | --- | --- |
| Association test | Number of methylation sites |  |  |
| $P$-value | All | Hypomethylated in NT1 (%) | |
| 1.00E-05 | 4 | 4 (100%) | 7.47E-01 |
| 0.0001 | 23 | 21 (91.3%) | 2.53E-01 |
| 0.001 | 147 | 123 (83.7%) | 1.55E-02 |
| 0.01 | 1,116 | 897 (80.4%) | 5.44E-09 |
| 0.05 | 4,650 | 3,749 (80.6%) | 1.73E-33 |
| Analyzed CpG sites*** |  |  |  |
|  | 735,841 | 455,080 (61.8%) |  |

\* Replicated probes only include probes for which the direction of methylation was consistent in both the discovery set and the replication set.
\*\* Hypothesis testing for the difference in the population proportions.
\*\*\* Probes located in the *HLA* region were excluded.

**Table 4.** NT1-associated hypomethylation sites are abundant in solo-WCGW.

|  | Rate in all<br>analyzed<br>CpGs | CD4<sup>+</sup> T cells |  |  |  |  | CD8<sup>+</sup> T cells |  |  |  |  |
| --- | --- | --- | --- | --- | --- | --- | --- | --- | --- | --- | --- |
|  |  | Rate in<br>hypomethylated<br>NT1 CpGs | *P*-value | OR | 95%CI |  | Rate in<br>hypomethylated<br>NT1 CpGs | *P*-value | OR | 95%CI |  |
|  |  |  |  |  | Lower | Upper |  |  |  | Lower | Upper |
| Solo-WCGW | 0.11 | 0.27 | 9.87E-194 | 3.069 | 2.839 | 3.317 | 0.21 | 1.46E-22 | 2.199 | 1.872 | 2.584 |
| Solo | 0.34 | 0.55 | 1.12E-150 | 2.446 | 2.283 | 2.622 | 0.52 | 9.64E-32 | 2.154 | 1.89 | 2.456 |
| WCGW | 0.23 | 0.45 | 2.85E-198 | 2.77 | 2.585 | 2.969 | 0.38 | 1.89E-29 | 2.136 | 1.867 | 2.444 |

Abbreviation: OR, odds ratio

To investigate the relationships between NT1-associated hypomethylated sites, we generated heatmaps of the methylation rates for the hypomethylated sites in each sample.

The analysis revealed a consistent pattern: individuals with a higher degree of hypomethylation showed this trait across almost all of these methylation sites, while those with a lower degree of hypomethylation displayed higher methylation across the board (Fig. S6). Consequently, the average methylation rate of these hypomethylated sites was used as an indicator to represent the degree of hypomethylation in each sample. We investigated the association between the degree of hypomethylation for each sample and clinical information, including age, sex, and body mass index (BMI). Sex and BMI did not show an association with the average value of hypomethylation in the discovery set (Fig. S7A, B). In the replication set, the BMI appeared to be weakly correlated with the average value of hypomethylation (R^2^ = 0.396); this may be attributable to the fact that the BMI was significantly higher in the NT1 cases than in the controls in the replication set. Age was weakly correlated with the average value of hypomethylation, and a tendency was observed for increased hypomethylation with higher age (R^2^ = 0.123-0.467; Fig. S7C). However, since age alone cannot explain the hypomethylation seen in NT1, we next examined the relationship between hypomethylation and various indicators that could be calculated with genome-wide methylation data using the DNA Methylation Age Calculator ^40^ and MethylDetectR ^41,42^ (Table S11). The analysis was adjusted for age. It should be noted that some of these indicators are based on values calculated assuming the input data is from whole blood, but our methylation data was obtained from fractionated T cells; thus, the reliability of these estimates was verified in the following section using whole blood methylation analysis data that we previously obtained with the 27K array. The analysis results showed that in the CD4^+^ T cells, granzyme A (GZMA) demonstrated the strongest association with hypomethylation (*P* = 2.26E-10), followed by correlations with hypoSC (a methylation-based mitotic index; *P* = 6.08E-06; Fig. 3D, E and Table S12). In CD8^+^ T cells, hypoSC showed the strongest association with hypomethylation (*P* = 1.61E-12), followed by a correlation with GZMA (*P* = 3.44E-08; Fig. 3D, E and Table S13).

### Investigation of NT1-related hypomethylation in whole blood

As described in the previous section, when exploring the indicators showing an association with hypomethylation in NT1 in the CD4^+^ and CD8^+^ T cells, some of the indicators were based on the assumption that the input data for estimation are methylation data derived from whole blood; thus, we also conducted studies using methylation data obtained from whole blood. We had previously reported that many NT1-related methylation sites were hypomethylated in NT1 when using whole blood for the analysis^28^. In the present study, to examine other details more thoroughly, we reanalyzed the data.

As shown in Fig. S8A, the *P*-values of the methylation sites hypomethylated in NT1 were significantly inflated when compared to the expected values. In a manner similar to our analysis of T cells, we then examined the relationship between the average methylation rates of hypomethylated sites associated with NT1 and the indicators calculated using the DNA Methylation Age Calculator ^40^ and MethylDetectR ^41,42^ to confirm the reproducibility of the results in the T cells. In our analysis, we focused particularly on hypoSC and GZMA. As a result, no significant association with the hypomethylation rate was observed when only age was adjusted for (Tables S12, S13 and Fig. S8B, C). However, upon reviewing the plots and adjusting for the effect of NT1, nominally significant associations between hypomethylation in NT1 and both hypoSC and GZMA were observed (hypoSC: *P* = 1.14E-4 and GZMA: *P* = 1.56E-2; Table S14 and Fig. S8D, E).

### Prevalence of solo-WCGW CpGs in NT1-associated hypomethylated sites

The hypoSC, an indicator related to cell mitosis, was among the indicators that showed an association with hypomethylation at NT1-related methylation sites. Previous studies have reported that DNA methylation at CpG sites is more likely to be lost in regions where there are fewer neighboring CpG sites, particularly in CpGs within the “WCGW” context (where W represents an A or T base; Solo-WCGW) ^43^. Following the previous study, we investigated the relationship between NT1-related hypomethylated sites and the number of CpG within windows of ±35 bp as well as the sequence context, including the bases on either side of the CpG (Table S15). We found that in both types of T cells, NT1-associated hypomethylated CpG sites were significantly more prevalent in solo-WCGW (CD4^+^ T cells: *P* = 9.87E-194, OR = 3.07 and CD8^+^ T cells: *P* = 1.46E-22, OR = 2.20; Table 4). Furthermore, upon detailed examination, it was found that NT1-associated hypomethylated sites were particularly prevalent within CpG sites located in the sequence of ACGT (Fig. S9).

## Discussion

In this study, we conducted an EWAS in CD4^+^ and CD8^+^ T cells from patients with NT1, and found that in both types of T cells, methylation sites associated with chemokine-related genes, such as *CCR4* and *CCL5*, were related to NT1. Particularly in CD4^+^ T cells, genes with expression levels that correlate with the methylation sites were predominantly involved in pathways of immune responses and cell chemotaxis. Additionally, we discovered the global hypomethylation of NT1-associated methylation sites in both cell types, and that the degree of hypomethylation was well-correlated with the estimated values of GZMA and hypoSC, indicators of cell mitosis. NT1-associated hypomethylated sites were significantly more prevalent in solo-WCGW context, which is known to lose methylation modifications easily during cell division, suggesting that cell division might be enhanced in the T cells of NT1 patients.

When we searched for methylation sites associated with NT1, to minimize false positives, we adopted two strategies. Firstly, instead of focusing only on single-point associations, *i.e.,* DMPs, we focused on DMRs. Secondly, even for single-point DMP associations, we sought to identify the DMPs that have been indicated to be associated with NT1 by other omics data. In the analysis for DMR identification, more DMRs were detected in the analysis of CD4^+^ T cells than in the analysis of CD8^+^ T cells (Table 1). However, most DMRs were found to be commonly associated in both cell types when including those that showed an association in either the discovery or replication analysis. Among these, especially in the replicated DMRs of both cell types, there were DMRs near *CCL5* and DMRs affecting the gene expression of *CCR4*. Both of these genes are chemokine-related, and considering that the previous GWAS of NT1 reported associations with a SNP, *i.e.,* rs3181077, located between other chemokine-related genes, *i.e., CCR1* and *CCR3*, and considering that the observed decrease in their expression levels in NT1, *CCL5* and *CCR4* represent highly intriguing candidate genes. *CCL5*, which is also known as regulated on activation, normal T cell expressed and secreted (RANTES), is a ligand for *CCR1*, *CCR3,* and *CCR5*, and it acts as a chemoattractant for monocytes, memory T-helper cells, and eosinophils ^44^. In this study, we were unable to find an association between the upstream DMR of *CCL5* and the gene expression level of *CCL5* itself, which was likely due to the very low expression levels of *CCL5* in the analyzed data. An investigation into the expression in immune cells using DICE (the database of immune cell expression, expression quantitative trait loci (eQTLs), and epigenomics) ^45^ revealed that the average transcripts per million (TPM) was less than 1 in all cell types. However, the single-cell analysis results showed expression primarily in CD4^+^ T cells ^45^. If we consider that the expression levels of genes with low expression levels, or in cases where *CCL5* expression levels are increased by some kind of immune system activation, there is a reasonable possibility of an association between gene expression and the NT1-associated DMRs in the *CCL5* region identified in this study. *CCR4* is a chemokine receptor for TARC/CCL17 and MDC/STCP-1/CCL22, which are mainly expressed by T cells ^46^. *CCR4* plays a crucial role in regulating immune balance, and is highly expressed in activated regulatory T cells; thus, it is increasingly being recognized as a potential target for cancer therapy ^46,47^.

The involvement of chemotaxis in NT1 was also suggested from the observed association with *S100A4* in the results of the integrated analysis of mQTLs using genotype data of CD4^+^ T cells (Table S5). *S100A4* is a member of the Ca(2+)-dependent S100 protein family and a transcription factor that regulates cellular motility. It is known that *S100A4* promotes filament depolymerization and chemotaxis through its direct interaction with soluble myosin-IIA, leading to the formation of stable protrusions ^48^. The gene expression of *S100A4* showed a negative correlation with methylation sites that were hypomethylated in the NT1 patient group, suggesting the possibility that, at least at the onset of NT1, the expression of *S100A4* was increased in the CD4^+^ T cells of the patients with NT1. A common issue in normal DNA methylation analysis is the difficulty in determining whether the associated factors are causes or consequences of the disease. In our analysis, it was also challenging to distinguish whether the associations found with many methylation sites were causes or consequences of NT1; however, the results of the integrated analysis with genotype data indicated causal effects, because genetic factors can be considered as causal for the disease. Therefore, it is highly likely that changes in the methylation rates affected by genetic variants (mQTLs) contributed to the disease onset. Considering this in conjunction with previous reports suggesting associations with chemokines derived from GWAS SNP associations ^12^, chemokine-related pathways could be a significant cause of NT1 development. Further detailed examinations of the chemokine pathways are necessary.

In this study, we observed global hypomethylation at NT1-related methylation sites in both CD4^+^ and CD8^+^ T cells (Fig. 2 and Tables 2, 3). This trend was not only seen in the replication set, but was also confirmed in the results of the methylation analysis for NT1 using whole blood that we had conducted previously ^49^. In our investigation of hypomethylated sites, we found that they were particularly scarce in CpG islands, and that they became more prevalent further from the islands. It was also evident that they were significantly less common in gene promoter regions and intragenic areas (Fig. 3). Although the methylation of CpG islands in the upstream promoter regions of genes greatly influences gene expression, much remains unknown about the functions of the methylation sites in such outside regions. While exploring why such hypomethylation occurs and what factors are involved, we discovered that the degree of hypomethylation was highly correlated with the levels of GZMA estimated from methylation data, as well as with the hypoSC cell division index (Fig. 3 and Tables S12, S13). We confirmed that there was very little overlap between the methylation sites used to calculate the degree of hypomethylation (CD4^+^ T cells: N = 3,259 and CD8^+^ T cells: N = 897) and those used to estimate indices, such as GZMA and hypoSC (N = 3,629), with only five sites in CD4^+^ T cells and one site in CD8^+^ T cells that overlapped. GZMA is one of the most abundant proteases present in the cytoplasmic granules of cytotoxic T cells and NK cells, and it is known for its crucial role in inducing the death of infected cells ^50^. However, in recent years, there has been debate regarding its role in cytotoxicity, and it has been shown that extracellular GZMA is involved in the regulation of inflammation and inflammatory diseases ^50^. The potential of GZMA to exacerbate inflammation and cause excessive immune responses, sepsis, tumor promotion, and rheumatoid arthritis has been suggested^50^. In the present study, the association of NT1 with cytotoxic granule exocytosis and inflammation was also suggested by the association of *NKG7* identified through the integrated analysis of methylation and gene expression data (Table S2). NK cell granule protein 7 is a factor that controls lymphocyte granule exocytosis and subsequent inflammation in a wide range of diseases, and it has been reported that *NKG7*, which is particularly expressed in CD4^+^ and CD8^+^ T cells, plays a crucial role in promoting inflammation in infections ^51^. Although determining whether the greater impact on hypomethylation stems from events like inflammation, associated with GZMA, or from the promotion of cell division, associated with hypoSC, is challenging due to the high correlation between their estimated values (CD4^+^ T cells: R^2^ = 0.528 and CD8^+^ T cells: R^2^ = 0.706), it is entirely plausible that T cell proliferation progressed due to the accelerated inflammation ^52^. The possibility of increased hypomethylation as a result of increased cell division was supported by the observation that NT1-related hypomethylated sites were especially prevalent in solo-WCGW motifs (Table 4). Previous studies have shown that hypomethylation occurs in only about half of the genome, especially in CG sequences not accompanied by surrounding CpGs and flanked by A or T bases, such as in WCGW motifs ^43^. It is also known that hypomethylation is more likely to occur in areas with low gene density, in areas with low GC density, and in so-called compartment B regions ^43,53^. These genomic characteristics overlap well with the features of regions where NT1-related hypomethylated sites were predominantly found. In other autoimmune diseases, particularly in systemic lupus erythematosus (SLE) and multiple sclerosis, global hypomethylation in immune cells has been reported ^54,55^. In the case of SLE, there have been reports suggesting that hypomethylated sites are often found upstream of immune-related genes, indicating that the mechanism may differ slightly from that of the hypomethylation we observed in NT1 ^54^. In multiple sclerosis, the global hypomethylation sites were particularly numerous within promoter regions in B cells, and they were reported to influence the expression of genes involved in the activation of B cells, which in turn further activates B cells ^55^. In contrast, the hypomethylation observed in NT1 was thought to possibly result from the progression of T cell division. Therefore, even though global hypomethylation is observed across different immune-related disorders, the mechanisms involved appear to be different. There are several limitations to this study. First, we obtained gene expression data from the same samples from which we acquired methylation data and conducted an integrated analysis, but the expression data was derived from the mRNA in whole blood.

If we had more samples available, it would have been preferable to obtain gene expression data from separated CD4^+^ and CD8^+^ T cells. Thus, we might have missed associations for some genes, especially those with varying expression patterns in different immune cells. Second, among the indicators showing a relationship with global hypomethylation in NT1, the estimated values for proteins, including GZMA, were based on predictions intended for methylation data derived from whole blood, but our methylation data was obtained from isolated T cells, leaving questions about the accuracy of these estimates. Although we have validated the factors of interest using methylation data from whole blood, this data was obtained using the 27K array, which has fewer probes. Therefore, it is desirable to confirm the reproducibility in the future using EPIC array data derived from whole blood. Thirdly, although we conducted a comprehensive methylation analysis using arrays, we cannot deny the possibility that methylation sites not covered by the arrays may be associated with NT1. In the future, when whole-genome bisulfite sequencing with a deeper depth than the sequencing depths possible now becomes available, more associated factors may be identified. Lastly, in this study, we first isolated CD8^+^ T cells from the buffy coat using magnetic beads, and then separated CD4^+^ T cells from the remaining blood components, resulting in the possibility that the CD8^+^ T cells included CD4^+^CD8^+^ double-positive T cells. Furthermore, we did not examine the different subtypes of CD4^+^ and CD8^+^ T cells. The global hypomethylation associated with NT1 showed a high correlation with indicators, such as GZMA and hypoSC, suggesting that inflammation and increased cell division could explain much of the hypomethylation. However, there were still differences in the degree of hypomethylation between the case and control groups that could not be fully explained. Namely, when comparing the same hypoSC and GZMA values, the NT1 cases tended to show more hypomethylation than the controls, particularly in the results of the CD8^+^ T cells and in the replication analysis with the 27K array using whole blood data. These data suggest the possibility of hypomethylation in NT1 that cannot be explained by inflammation or increased cell division alone. Particularly, the significant fact that hypomethylation cannot be explained by GZMA and hypoSC alone in the whole blood data indicates that the presence of different subtypes of immune cells in NT1 compared to control individuals may be a contributing factor. Future single-cell analyses in NT1 could potentially reveal the details of the immune abnormalities that contribute to the onset of the disease.

In conclusion, the results of our DNA methylation analysis of CD4^+^ and CD8^+^ T cells in NT1 suggest a high likelihood that abnormalities in chemokine-related pathways are a major cause of disease onset. Furthermore, our results indicate that increased T cell division in NT1, possibly driven by heightened inflammatory responses, may lead to hypomethylation in regions of DNA where methylation is more easily disrupted. Future investigations, including single-cell analyses and single-cell DNA methylation analyses, are expected to reveal more details of the mechanism of NT1 onset.

## Methods

### Subjects in the discovery and replication sets

Our research design is shown in Fig. 1. Samples from 42 (discovery: 28, replication: 14) patients with NT1 and 42 (discovery: 28, replication: 14) control subjects without significant sleep-disturbing events who underwent diagnostic sleep studies (polysomnogram and the multiple sleep latency test) at Seiwa Hospital (Shinjuku, Tokyo, Japan) or Koishikawa Tokyo Hospital (Bunkyo, Tokyo, Japan) were included. All subjects were Japanese. The demographic characteristics are shown in Table S16 and S17. NT1 was diagnosed by sleep specialists according to the International Classification of Sleep Disorders, 3rd edition ^56^, and all patients carried the *HLA-DQB1*06:02* allele. Fasting blood was drawn at 7:00 am after a polysomnogram, and all subjects were drug-naïve or had stopped medication that could affect sleep at least 1 week before the time of blood collection. We used two EDTA-2Na-coated blood collection tubes: one for buffy coat DNA extraction and the other for lymphocyte separation by density-gradient centrifugation using Ficoll-Paque Plus (Merck Millipore, USA). The lymphocyte fraction was washed twice with phosphate-buffered saline, resuspended in Cellbanker 2 (Zenogen Pharma, Japan), and stored at −80°C until use. We simultaneously collected blood into two Paxgene RNA collection tubes (Beckton-Dickinson, Franklin Lakes, NJ, USA), and extracted RNA using a Paxgene Blood RNA kit (QIAGEN, Hilden, Germany) according to the manufacturers’ protocols. The extracted RNA was used for RNA-seq as described below. This study received ethical approval from all relevant agencies, and written informed consent was obtained from all participants.

### Isolation of CD4^+^ and CD8^+^ T cells and DNA extraction

CD4^+^ and CD8^+^ T cells were extracted from the lymphocyte fraction using the Dynabeads CD4 Positive Isolation Kit and Dynabeads CD8 Positive Isolation Kit (Invitrogen, Waltham, MA, USA) according to the manufacturer’s protocols. First, CD8^+^ T cells were isolated from the buffy coat, and CD4^+^ T cells were isolated from the remaining blood cell components. DNA was extracted from both types of T cells using the QIAamp DNA Mini Kit (QIAGEN).

### DNA methylation analysis

DNA was first bisulfite-converted, then the genome-wide methylation profile was examined with a DNA methylation array (EPIC array; Infinium® Methylation EPIC BeadChip, Illumina). The methylation arrays were imaged using a high-precision scanner (iScan System, Illumina); the scan data (idat files) were captured with the ChAMP R package ^57–59^. The methylation rate was calculated as the β value and the M values (the logit transformations of the β values) were used in the case-control analysis. Detection P values were calculated using a method reported by Heiss *et al.* ^60^.

### Filtering and normalization of the methylation data

Poor-performing probes were excluded from the analysis according to a method reported by Zhou *et al.* (Fig. S1, S10) ^61^. Data normalization involved three steps: (1) correcting the bias between Infinium I and II assay systems using beta mixture quantile dilation ^62^, (2) applying quantile normalization to adjust for variations in methylation rate distributions between samples, and (3) using ComBat ^63^ with T cell type (CD4^+^ or CD8^+^) as a covariate to correct for batch effects.

### Quality control of the methylation data

The following three analyses were performed for quality control:

1. PCA using methylation data for all sites to check for bias in the distribution of the PCA plot between case and control samples. In a previous study, when PCA was performed using genome-wide methylation rate data for CD4^+^ T cells, the principal components were reported to be mainly explained by the methylation rate of the GATA binding protein 3 (GATA3) binding sites ^32^. GATA3 is the master transcription factor of T cells, and it controls the differentiation and function of each T cell subset ^64^. Therefore, we additionally checked whether the methylation rate of the binding sites of master transcription factors, *i.e.,* T-bet and GATA3, explained the main component in our PCA as well. We utilized previously reported ChIP-Seq data for T-bet and GATA3 ^65^, and for the methylation rate of each master transcription factors’ binding site, dimensionality reduction by PCA was performed and PC1 was used as a feature.
2. Estimation of the cellular composition within each sample. In this study, although we separated CD4^+^ and CD8^+^ T cells for the methylation analysis, there are subsets of each cell type that can be further subdivided. To confirm whether the composition of such subsets differed between the case and control groups, we estimated the cellular composition using MeDeCom, which does not require any reference methylation data ^66^. In addition, we also performed reference-based cellular composition estimation as reported by Horvath ^40^ to estimate the percentage of CD4^+^ T cells and CD8^+^ T cells in the samples to confirm that each sample was properly estimated to contain mainly CD4^+^ T cells or CD8^+^ T cells. Samples in which the most abundant cell type was not correctly estimated to be the expected CD4^+^ T cells or CD8^+^ T cells were excluded from the subsequent association analyses.
3. Estimation of age using methylation data. The biological age can be estimated from methylation data ^40^. We examined whether the biological age was accelerated compared to the chronological age in NT1 patients, and we also examined whether the ages of the controls were properly estimated.

### Association analysis of the methylation data

To detect NT1-associated methylation sites, we searched for DMPs and DMRs using the *t* test and bump hunting method ^67^, respectively. For DMPs, since not only the statistical significance from P values, but also a large difference in the methylation rate is important for biological significance, we calculated a combined rank considering both indices (the *P* value and Δ*β,* which is the average *β* for the cases – average *β* for the controls), and we considered the top-ranked NT1-associated methylation sites to be particularly important. Association analyses and visualization of the data, such as Q-Q plots, were conducted using R software (ver. 3.6.2).

### RNA sequencing

We extracted RNA from the whole blood samples of the 42 patients with NT1 and 42 control subjects included in the discovery and replication sets for DNA methylation analysis and performed RNA-seq. Sequencing libraries were prepared using TruSeq Stranded Total RNA Ribo Zero (Illumina), and paired-end RNA-seq was performed on a NovaSeq 6000 (Illumina). The FASTQ read data were trimmed using a quality score of 0.05 and a minimum length of 15 nt in CLC Genomics Workbench 21 (QIAGEN). Read mapping was performed with the following parameters: mismatch cost, 2; insertion cost, 3; deletion cost, 3; length fraction, 0.8; similarity fraction, 0.8; and maximum number of hits for a read, 10. The read count data were corrected by the trimmed mean of M values (TMM) method implemented in the edgeR package ^68–70^. Association analysis of the expression data was performed using the likelihood ratio test with age, sex, and BMI as covariates in the design matrix.

### Genotyping

DNA samples that were included in the DNA methylation analysis and RNA-seq (NT1: N = 42 and control: N = 42) were genotyped with Axiom Japonica array v2 (Toshiba, Tokyo, Japan). Imputation analysis was performed using BEAGLE5.1 ^71^ with reference panels from the 1000 Genomes Project (phase 3) and from the Tohoku Medical Megabank 14KJPN ^72^. SNP filtering was performed with the following criteria: information score ≥ 0.5; minor allele frequency ≥ 0.01; SNP call rate ≥ 95%; and Hardy-Weinberg equilibrium P value > 0.01. Genotype data from a previous GWAS of NT1 were also used in the analysis. The details of the GWAS data are described elsewhere ^12,73^.

### Integrative analysis of methylation and gene expression

To perform an integrative analysis of the methylation and gene expression data, we first detected pairs of methylation rate and gene expression that correlated with each other (Methylation-Expression pairs) using data from previous studies with larger sample sizes (Table S1) ^34–38^. Next, we searched for pairs in which the correlated methylation rate and gene expression were both associated with NT1 in the samples of this study, and in which the direction of the association was consistent with the Methylation-Expression pair (Fig. 1B). We searched for previous studies that measured genome-wide methylation rates and gene expression in the same sample of CD4^+^ T cells or CD8^+^ T cells, and had a sample size of N > 50. For CD4^+^ T cells, we included the three previous studies shown in Table S1, while for CD8^+^ T cells, there were no studies that reported EPIC (Illumina) or sequence-based methylation data, and only data from the 450K array (Illumina) were available. Since our primary focus was on CD4^+^ T cell analysis from the results of the association analysis of NT1, for the CD8^+^ T cells, we conducted an integrative analysis with the 450K data as a preliminary analysis. For the methylation data of the previous studies, we used filtered and normalized data, if available. For studies with only raw methylation data, standard filtering and normalization of ChAMP were performed. For expression data, we used the data processed in each study, and gene identifiers on each platform were converted to Ensembl identifiers using information from GENCODE ^74^ and BioMart ^75^.

Correlated Methylation-Expression pairs were explored using Matrix eQTL ^76^ with age and sex as covariates. In the analysis of Study 1, age was estimated from the genome-wide methylation data and used as a covariate, because no age information was available for Study 1 (Table S1). In Study 3 (Table S1), only sex was used as a covariate, because the subjects were children aged 0 to 2 years. We specifically focused on Methylation-Expression pairs in *cis* (<1 Mb), and identified pairs under two conditions: *P* < 0.05 and FDR < 0.05.

Among the detected Methylation-Expression pairs, we searched for pairs in which both the methylation rate and gene expression were related to NT1 (Fig. 1B). An NT1-associated methylation site, *i.e.,* a DMP, was defined as a site with a *P* < 0.05 and a combined rank within the top 1% among the discovery samples, and was replicated with a *P* < 0.05 in the replication samples. NT1-associated gene expression was defined as expression with a *P* < 0.05 and FC >25% using our RNA-seq data.

### Integrative analysis of methylation and genotype data

To integrate the genotype and methylation data, we used information from the mQTL. An mQTL is where the methylation rate correlates with the genotype of the SNP, and it has been reported that the mQTL profiles are nearly identical between whole blood and each immune cell ^77^. We searched for previous studies that used whole blood with a large sample size, and used the mQTL information reported in the two studies shown in Table S4 ^77,78^. Among the reported mQTLs, we searched for those that had a methylation rate and SNPs that were both associated with NT1, and whose direction of association was consistent with the previous studies. For SNPs, we focused on those that were associated with NT1 in our previous GWAS to exclude SNPs that showed associations only in this study due to the small sample size (NT1: N = 42 and control: N = 42), and not in the previous GWAS of NT1 with larger sample size ^12^ (Fig. 1C). The detailed conditions for the mQTL search were as follows: an NT1-associated methylation site, *i.e.,* a DMP, was defined as a site with a *P* < 0.05 and a combined rank within the top 1% among the discovery samples, and was replicated with a *P* < 0.05 in the replication samples. NT1-associated SNPs were defined as those with a *P* < 1E-3 in the GWAS of NT1, and in the present sample, when the OR > 1, those with an OR greater than the OR in the GWAS, and when the OR < 1, those with an OR smaller than the OR in the GWAS (Fig. 1C).

### Pathway analysis

In the CD4^+^ T cell analyses, since NT1-associated methylation sites were suggested in the analysis for DMRs and in the integrative analyses with gene expression data and genotype data, pathway analysis was conducted using genes whose expression was affected by the methylation sites. The gene expressions affected by the methylation sites were linked based on the relationship of Methylation-Expression pairs, as determined through the integrative analysis of gene expression data and methylation data. Pathway analysis was performed using MetaCore™ software (version 6.24 build 67895, Clarivate, Boston, MA, USA), and the PANTHER Classification System (PANTHER 17.0) ^79^ was used to validate the results from the MetaCore software. We then searched for significant gene sets that met the following conditions in the Gene Ontology database ^80,81^: i) FDR < 0.05 in the MetaCore analysis, and replicated in PANTHER with *P* < 0.05; ii) The pathway includes three or more NT1-related genes; iii) The total number of objects in the pathway is less than 3,000. The detected pathways were clustered using the k-means method based on the presence or absence of NT1-related genes associated with each pathway. The number of clusters, *k*, was set to four based on manual curation and intuitive grouping observed within the detected pathways.

### Investigation of NT1-related hypomethylated sites

Since a global hypomethylation trend was observed in NT1, we examined the NT1-associated hypomethylation sites with a *P* < 0.01 in the discovery set that were replicated with a *P* < 0.05. To explore the kinds of genomic regions where the hypomethylated sites are commonly present, Ensembl Human Regulatory Features (GRCh38.p14) was used to examine the positional relationship between hypomethylated sites and genomic regulatory features, such as enhancers and promoters ^82^. We extracted the data originating from T cells (Table S18). GENCODE (version 28) data were used to determine the positional relationship of regions related to genes ^74^. We also used the functional genomics data obtained from the ENCODE project ^83^ to investigate genomic features by Histone ChIP-seq. The experimental matrix was filtered by the following conditions: status, released; perturbation, not perturbed; organism, *Homo sapiens*; cell, T cell; available file types: bed narrowPeak; and all audit category filters were applied. The positional information for CpG islands was based on Illumina’s annotation file for the EPIC array. Since the genomic regulatory features from Gencode and Encode projects were based on hg38, the positional information of hypomethylated sites (hg19) was converted by the NCBI Genome Remapping Service for comparisons ^84^.

To visualize the relationships between hypomethylated sites, we generated heat maps, and confirmed that the methylation rates were generally low in samples with low levels of methylation in the NT1-associated hypomethylation sites. Since some subjects showed a generally high methylation status and others showed a generally low hypomethylation status across the selected DMPs, the average methylation rates of NT1-associated hypomethylation sites were calculated for each sample, serving as an indicator of the degree of hypomethylation. To identify factors that could explain this degree of hypomethylation, correlations with age, sex, and BMI were examined.

To examine factors associated with the degree of hypomethylation, we calculated various indicators that could be determined from genome-wide methylation data using the DNA Methylation Age Calculator ^40^ and MethylDetectR ^41,42^ (Table S11). We then conducted a linear regression analysis using age as a covariate to explore indices indicating an association with the degree of hypomethylation. Some indices were designed assuming that the methylation data were derived from whole blood as the input. Therefore, for the validation of the results, similar estimations were also conducted using data from an EWAS of whole blood from patients with NT1 that we had previously performed ^28^. The prior study included 26 cases of NT1 and 20 controls, and the methylation levels were measured using the 27K array (Illumina); the details of the analysis are described elsewhere ^28^.

## Acknowledgements

This study was supported by Grants-in-Aid for Scientific Research (Grant numbers 15H04709, 17J01616, 18K15053, 19H03588, 21H02856, and 22H03006) from the Ministry of Education, Culture, Sports, Science and Technology of Japan. This study was also supported by the Practical Research Project for Rare/Intractable Diseases and the Strategic Research Program for Brain Sciences from the Japan Agency for Medical Research and Development (AMED). We thank all participants who cooperated in this study. We also thank the staff of Seiwa Hospital and Koishikawa Tokyo Hospital for their support in blood sampling.

## Authors’ contributions

TM and MS designed the study. MH collected the blood samples. MS and YH performed the experiments. MS and TM carried out the statistical analyses. TM, YH, KT, TK and MH interpreted the results. MS, TM, and MH drafted the manuscript, and all authors contributed to the final version of the paper.

## Data availability

The genome-wide methylation dataset supporting the conclusions of this article has been deposited in the DNA Data Bank of Japan (DDBJ; https://www.ddbj.nig.ac.jp/indexe.html), and is available from the European Nucleotide Archive operated by the European Molecular Biology Laboratory’s European Bioinformatics Institute (EMBL-EBI; https://www.ebi.ac.uk/ena/browser/home) under the accession number PRJDB15980.

