## Supplementary Figures for "Multiomics analysis of narcolepsy T cells: global hypomethylation linked to T cell proliferation"

**Fig. S1.** Procedure for filtering out the poor-performing probes in the discovery set.
**Fig. S2.** Principal component analysis (PCA) using genome-wide methylation data.
**Fig. S3.** Estimation of the cellular composition ratios using MeDeCom in the discovery set.
**Fig. S4.** Comparison of the estimated cellular composition ratios between the case and control groups.
**Fig. S5.** Estimated DNA methylation age.
**Fig. S6.** Heatmap of the methylation rates at hypomethylated sites in NT1.
**Fig. S7.** Association between the degree of hypomethylation and clinical information.
**Fig. S8.** Re-examination of the EWAS data from the whole blood-derived DNA of NT1 patients analyzed with the 27K array.
**Fig. S9.** Examination of the sequences surrounding the analyzed CpG sites.
**Fig. S10.** Procedure for filtering out the poor-performing probes in the replication set.

The Supplementary Tables will be provided in a separate Excel file.

**Table S1.** Previous studies used for the integrative analysis of methylation and gene expression
**Table S2.** Results of the Methylation-Expression analysis in $CD4^+$ T cells
**Table S3.** Results of the Methylation-Expression analysis in $CD8^+$ T cells
**Table S4.** Previous studies used for the integrative analysis of mQTL
**Table S5.** Result of mQTL analysis in $CD4^+$ T cell
**Table S6.** Result of mQTL analysis in $CD8^+$ T cell
**Table S7.** Results of the pathway analysis with genes related to NT1-associated methylation sites in $CD4^+$ T cells
**Table S8.** The relationship between NT1-related hypomethylation sites and CpG islands
**Table S9.** The relationship between NT1-related hypomethylation sites and gene regions
**Table S10.** The relationship between NT1-related hypomethylation sites and regions marked by histone modifications
**Table S11.** Indicators calculated with the DNA Methylation Age Calculator and MethylDetectR
**Table S12.** Indicators associated with hypomethylation in the $CD4^+$ T cell analysis
**Table S13.** Indicators associated with hypomethylation in the $CD8^+$ T cell analysis
**Table S14.** HypoSC and GZMA associations in whole blood after adjusting for the effect of NT1

**Table S15.** Number of CpGs classified in the adjacent base context
**Table S16.** Demographic characteristics of the discovery set
**Table S17.** Demographic characteristics of the replication set
**Table S18.** Types of T cells registered in Ensembl Human Regulatory Features used for analysis

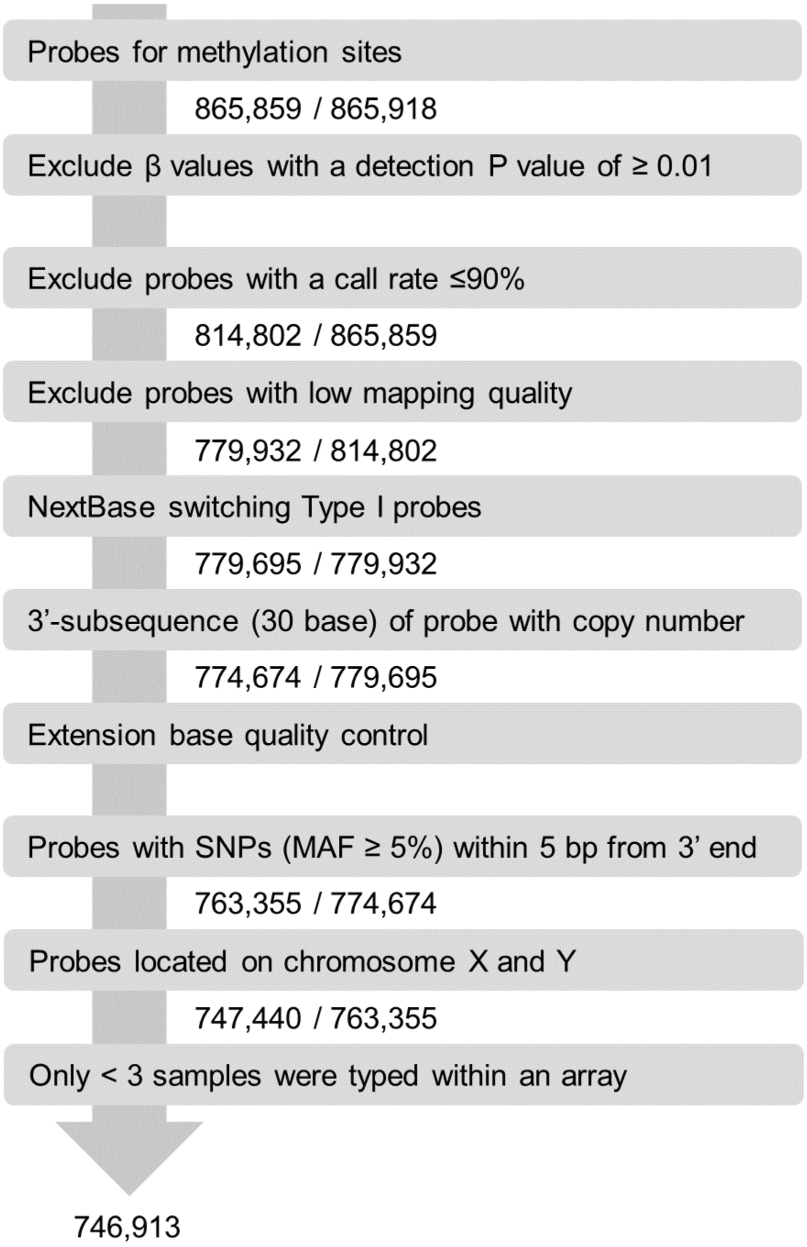


**Fig. S1.** Procedure for filtering out the poor-performing probes in the discovery set. First, for each sample, probes with a detection *P*-value greater than 0.01 were identified and taken to be unreliable for typing purposes. In the analysis of all samples, probes that failed to be typed in at least three samples were excluded. Furthermore, probes that have been reported to be poor-performing in previous studies, and probes on sex chromosomes were excluded.

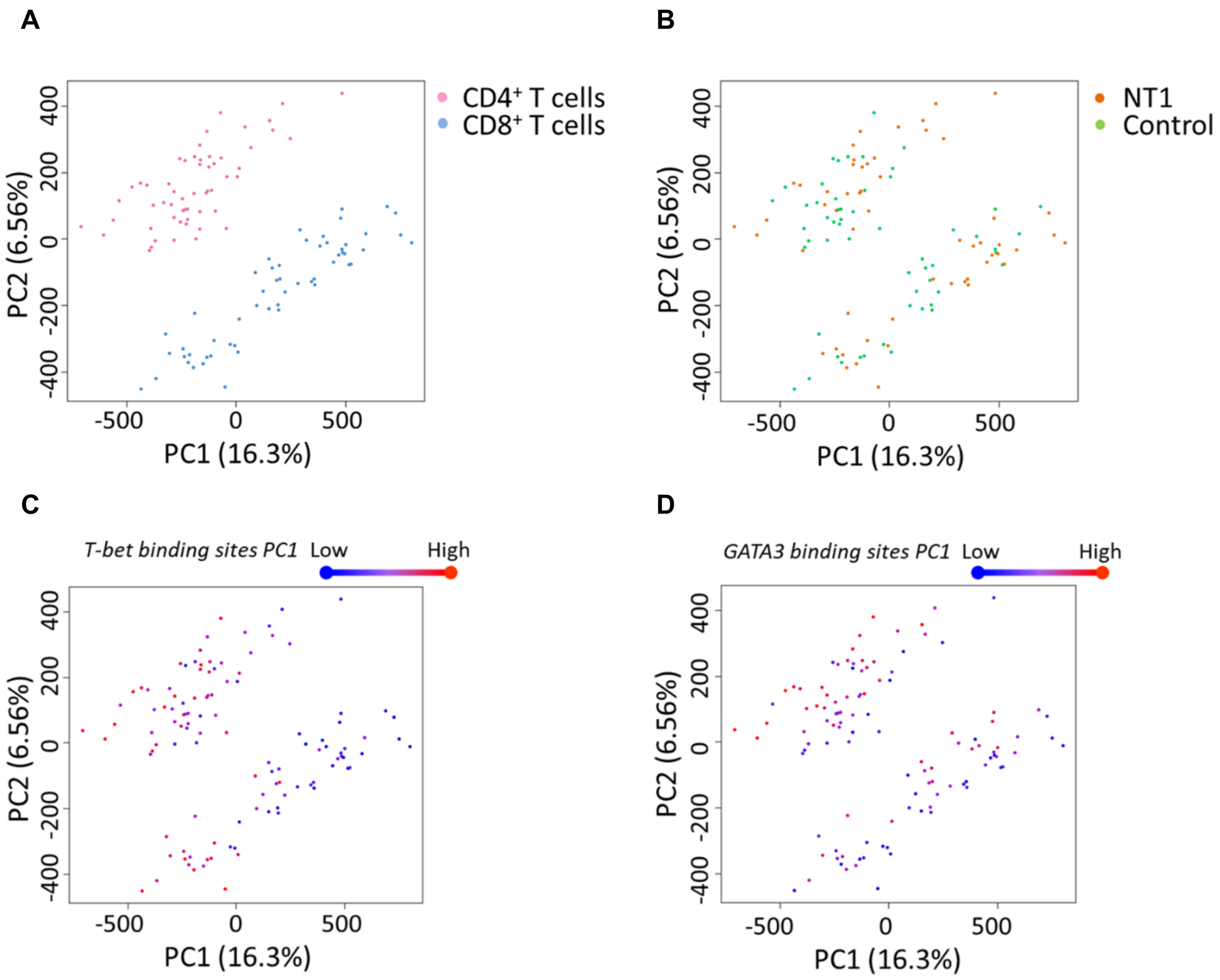


**Fig. S2. Principal component analysis (PCA) using genome-wide methylation data. A.** PCA with color differentiation for the two types of T cells. **B.** PCA with color differentiation for the NT1 and control groups. **C.** PCA with color differentiation for the methylation levels at T-bet binding sites. **D.** PCA with color differentiation for the methylation levels at GATA3 binding sites.

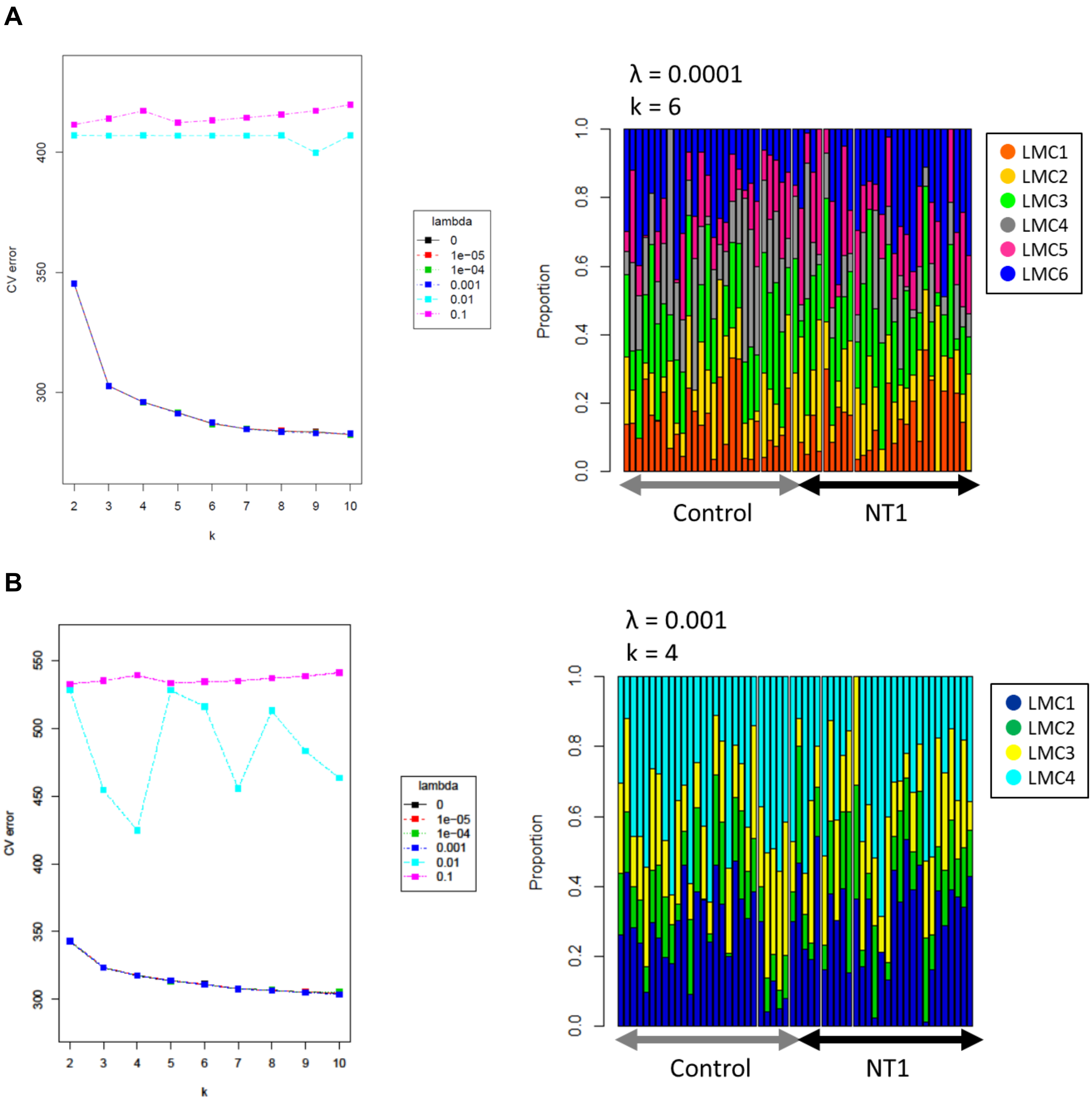


**Fig. S3. Estimation of the cellular composition ratios using MeDeCom in the discovery set.** In this study, we measured the methylation levels after fractionating the CD4$^+$ and CD8$^+$ T cells. Furthermore, to investigate the possibility that more refined subtypes may vary between the case and control groups, we used the genome-wide methylation data to estimate the cellular composition ratios using a reference-free approach called MeDeCom. **A.** Results for the CD4$^+$ T cells. With consideration to reducing the cross-validation (CV) error, we selected λ = 0.0001 and k = 6 as being optimal. **B.** Results of the CD8$^+$ T cells. With consideration to reducing the CV error, we selected λ = 0.001 and k = 4 as being optimal.

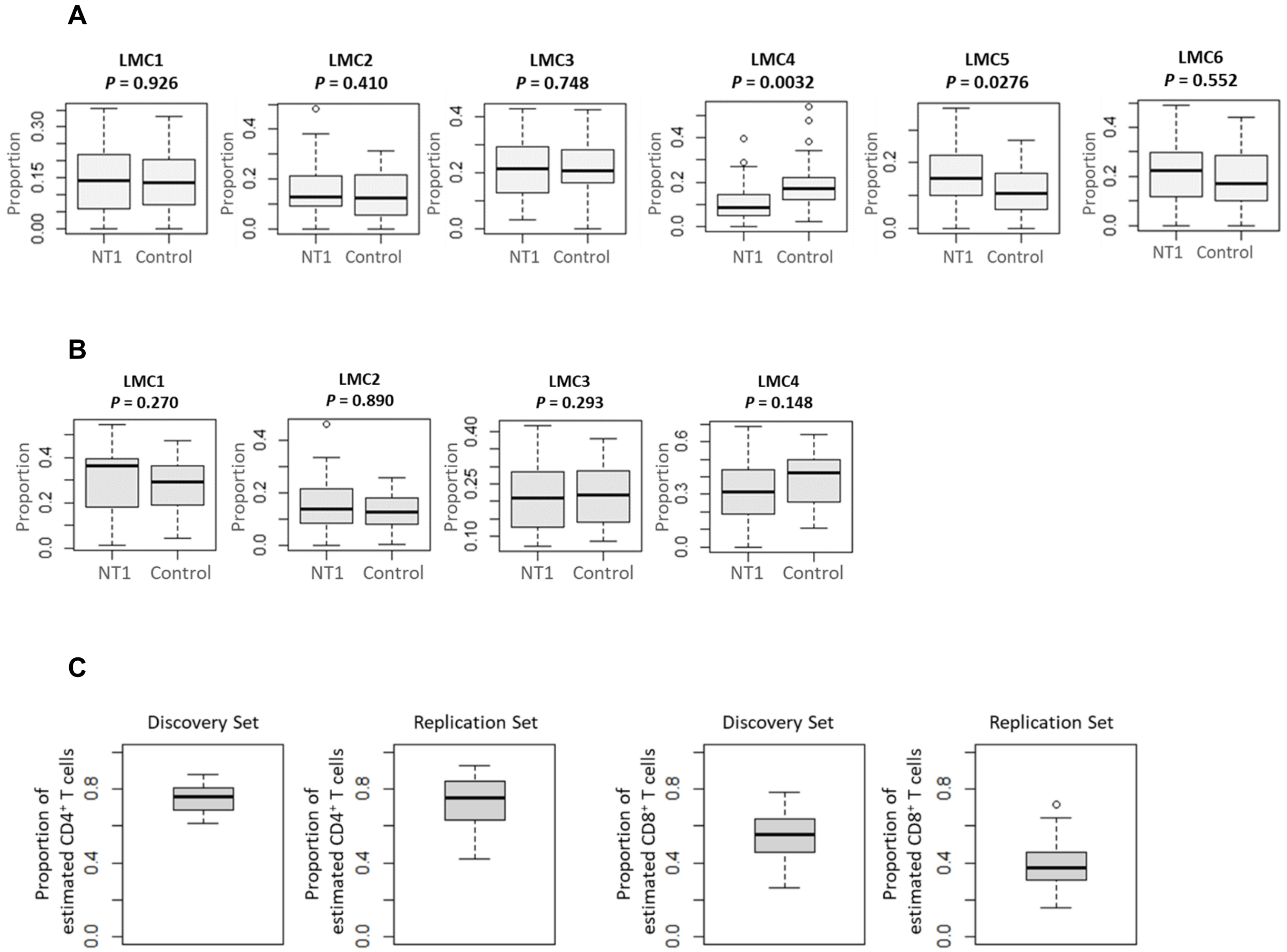


**Fig. S4. Comparison of the estimated cellular composition ratios between the case and control groups. A.** Case-control comparisons of the latent methylation components (LMCs) inferred from the MeDeCom analysis of the $CD4^+$ T cell data in the discovery set. **B.** Case-control comparisons of the LMCs inferred from the MeDeCom analysis of the $CD8^+$ T cell data in the discovery set. **C.** The proportion of cells classified as $CD4^+$ T cells in the $CD4^+$ T cell data, and the proportion of cells classified as $CD8^+$ T cells in the $CD8^+$ T cell data in the reference-based analysis.

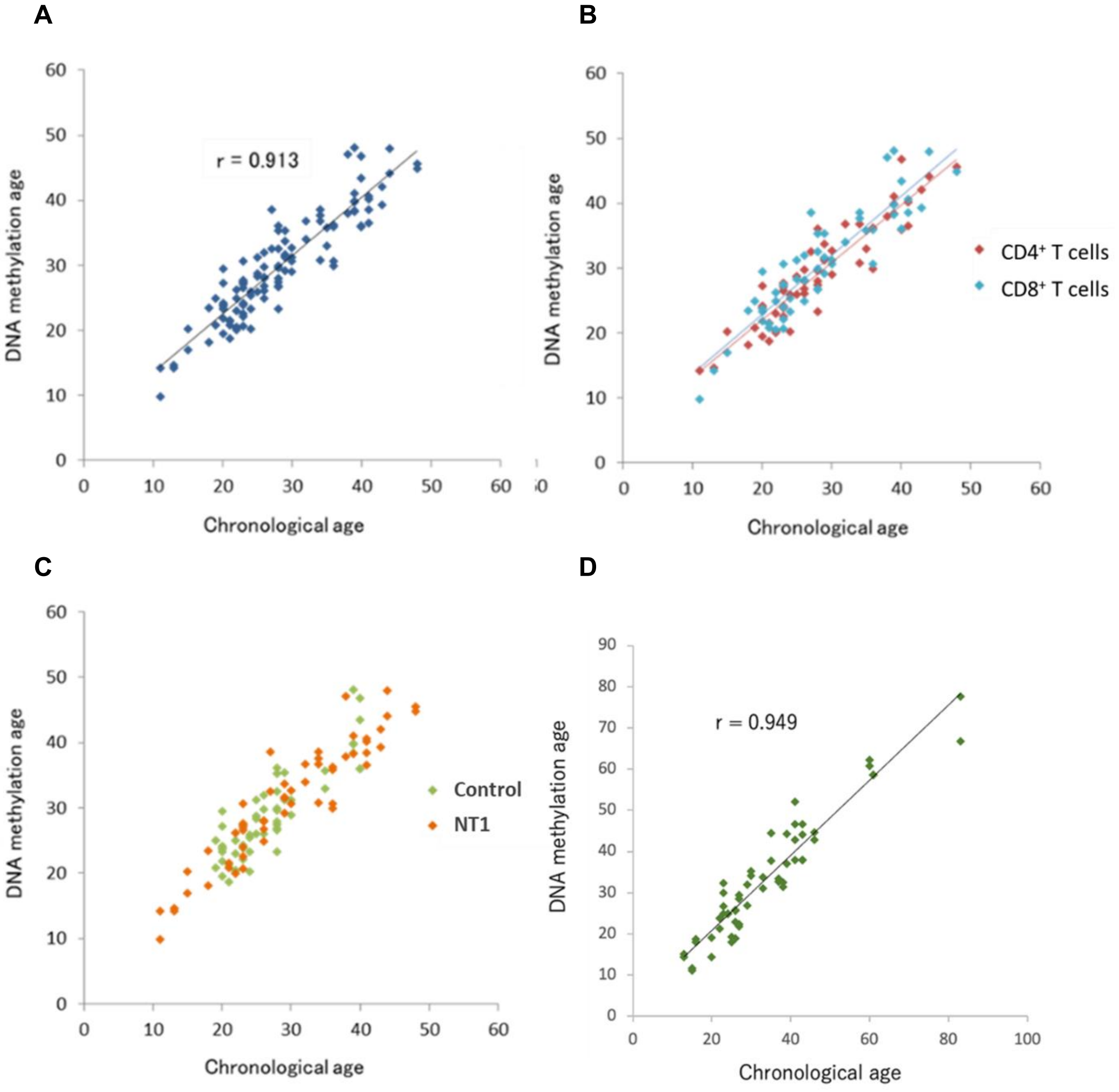


**Fig. S5. Estimated DNA methylation age. A.** Estimated DNA methylation age in the discovery set. A strong correlation was seen between the DNA methylation age and chronological age (correlation coefficient: 0.913). **B.** Comparison of the estimated methylation age between the $CD4^{+}$ and $CD8^{+}$ T cells in the discovery set. **C.** Comparison of the estimated methylation age between the case and control groups in the discovery set. **D.** Estimated DNA methylation age in the replication set. A strong correlation was seen between the DNA methylation age and chronological age (correlation coefficient: 0.949).

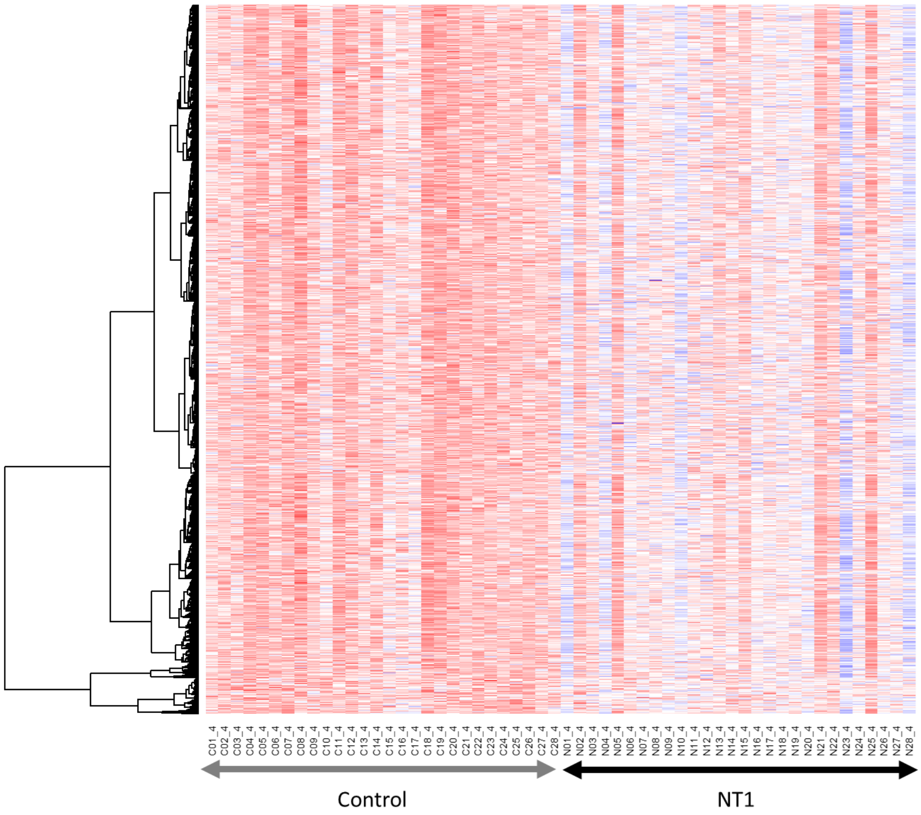


**Fig. S6. Heatmap of the methylation rates at hypomethylated sites in NT1.** We examined methylation sites that were associated with NT1 with a $P < 0.01$ in the discovery set, were replicated with a $P < 0.05$, and were hypomethylated in NT1. The vertical axis shows each of the NT1-associated hypomethylated sites, while the horizontal axis shows each of the samples. Across these methylation sites, it was observed that samples with a stronger overall degree of hypomethylation exhibited hypomethylation at most of the methylation sites while samples with a weaker degree of hypomethylation did not exhibit hypomethylation at most of the methylation sites.

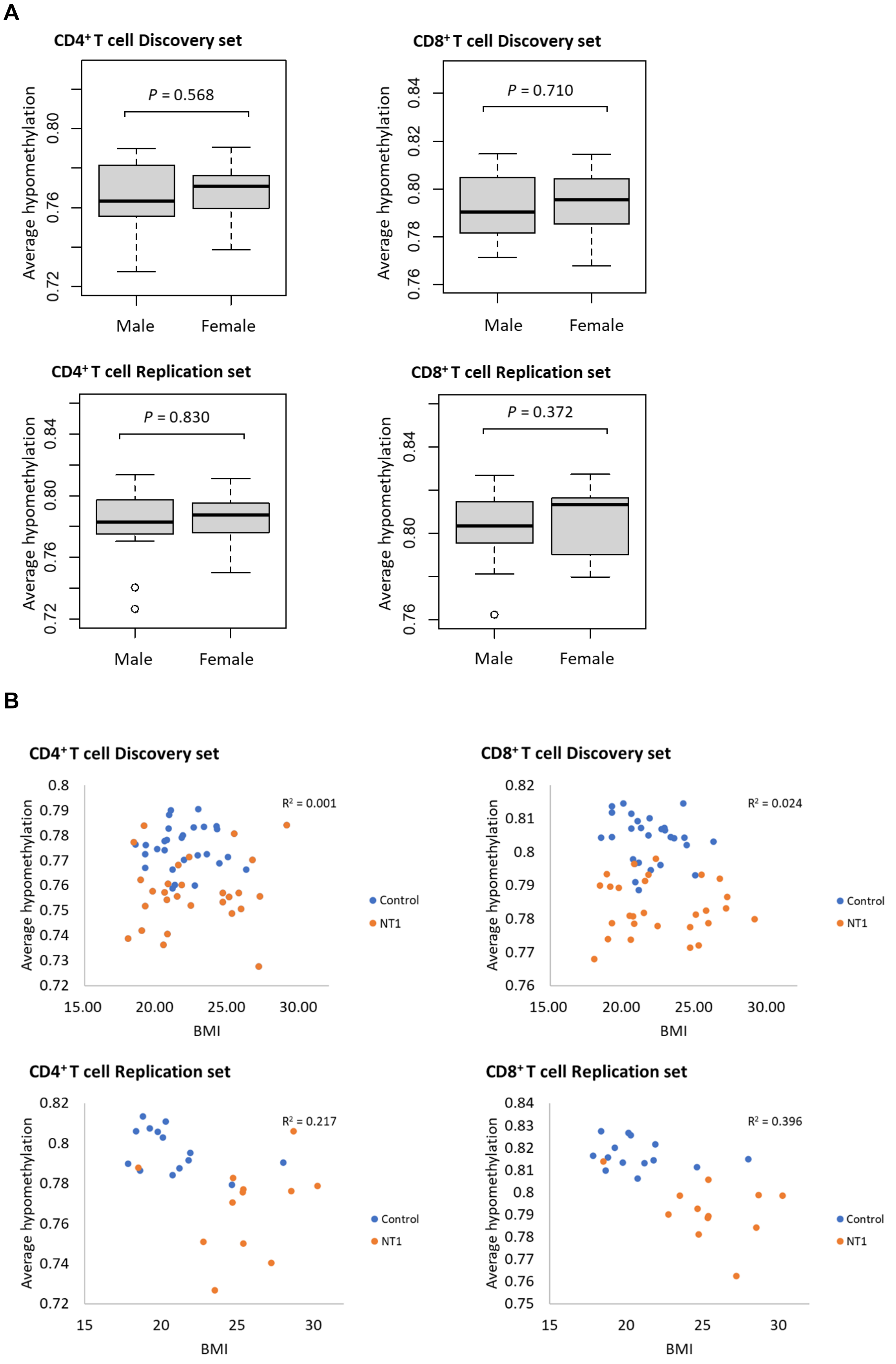

A
CD4⁺ T cell Discovery set
P = 0.568
Average hypomethylation
0.72
0.76
0.80
Male
Female
CD8⁺ T cell Discovery set
P = 0.710
Average hypomethylation
0.76
0.78
0.80
0.82
0.84
Male
Female
CD4⁺ T cell Replication set
P = 0.830
Average hypomethylation
0.72
0.76
0.80
0.84
Male
Female
CD8⁺ T cell Replication set
P = 0.372
Average hypomethylation
0.76
0.80
0.84
Male
Female
B
CD4⁺ T cell Discovery set
R² = 0.001
Average hypomethylation
0.8
0.79
0.78
0.77
0.76
0.75
0.74
0.73
0.72
15.00
20.00
25.00
30.00
BMI
Control
NT1
CD8⁺ T cell Discovery set
R² = 0.024
Average hypomethylation
0.82
0.81
0.8
0.79
0.78
0.77
0.76
15.00
20.00
25.00
30.00
BMI
Control
NT1
CD4⁺ T cell Replication set
R² = 0.217
Average hypomethylation
0.82
0.8
0.78
0.76
0.74
0.72
15
20
25
30
BMI
Control
NT1
CD8⁺ T cell Replication set
R² = 0.396
Average hypomethylation
0.84
0.83
0.82
0.81
0.8
0.79
0.78
0.77
0.76
0.75
15
20
25
30
BMI
Control
NT1

**C**

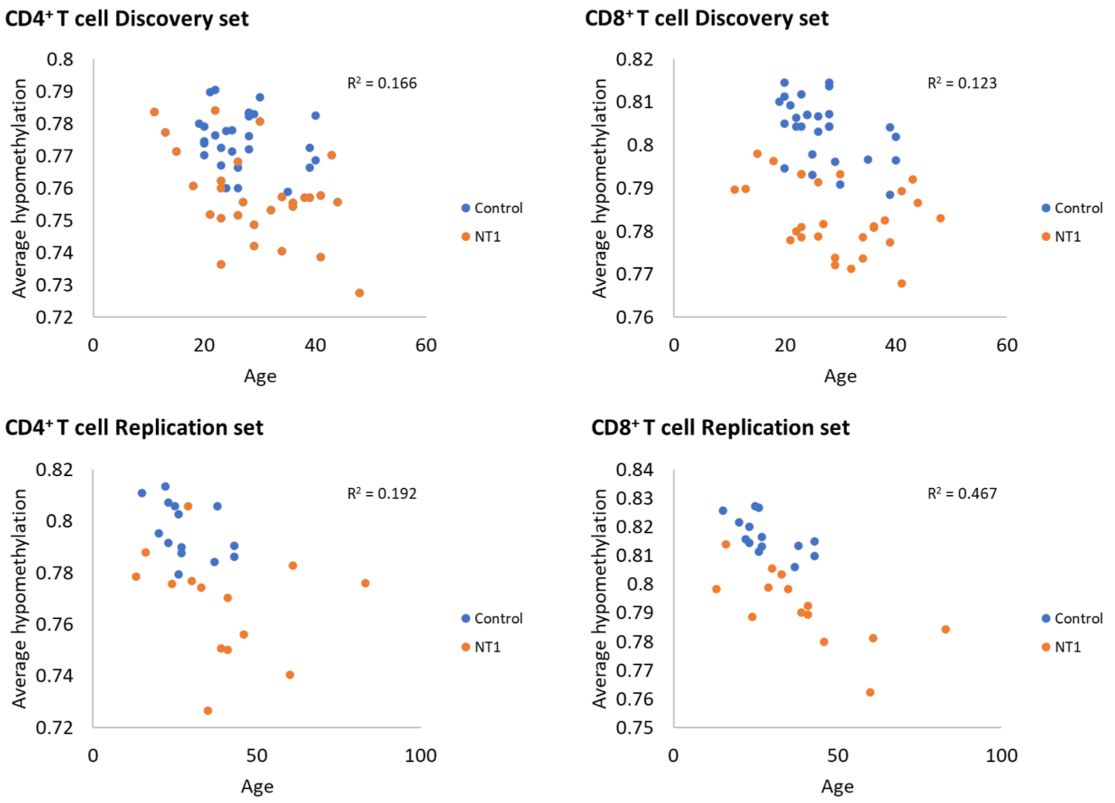


**Fig. S7. Association between the degree of hypomethylation and clinical information.** **A.** Association between the degree of hypomethylation (average hypomethylation) and sex. **B.** Association between the degree of hypomethylation and BMI. **C.** Association between the degree of hypomethylation and age.

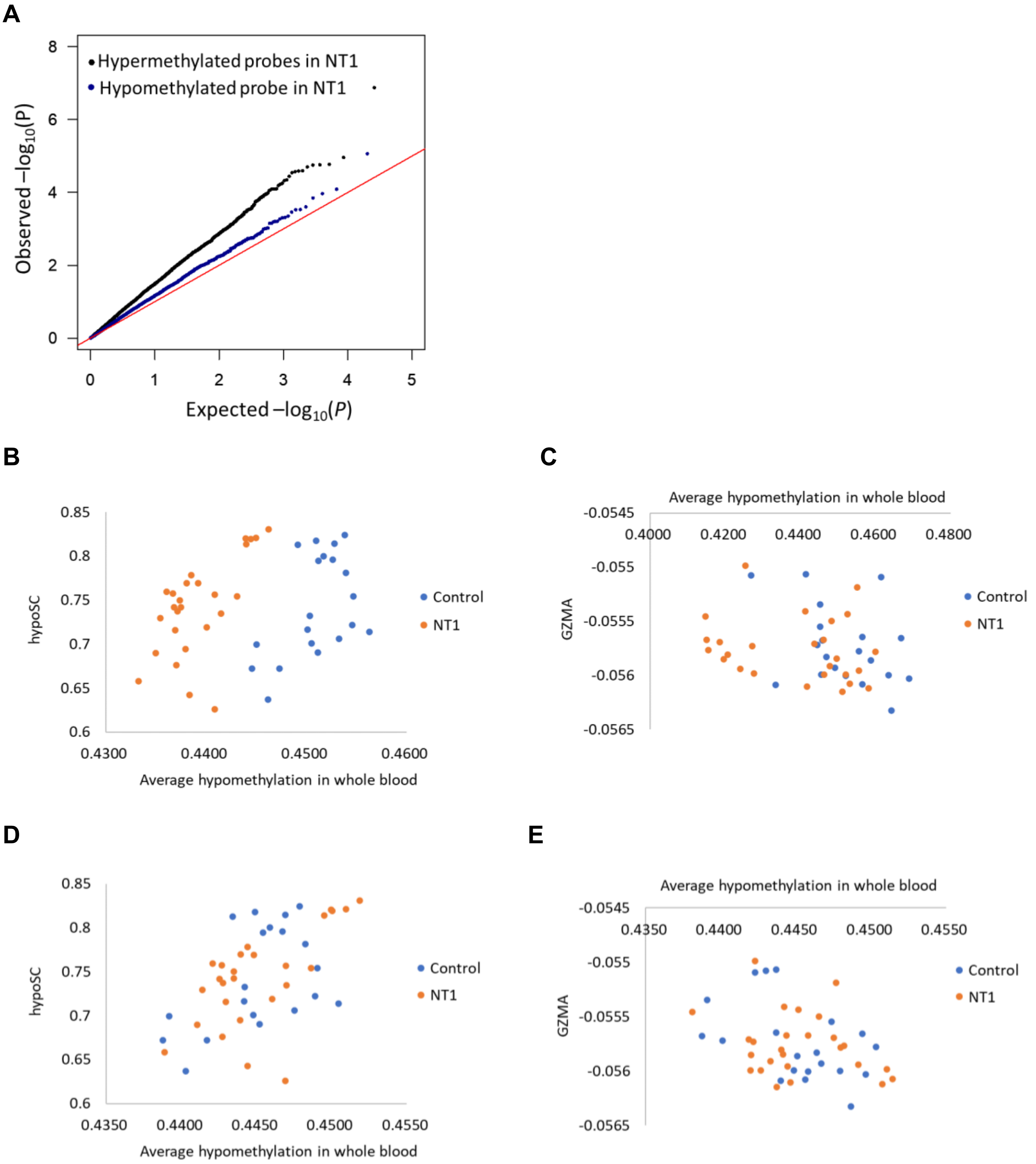


**Fig. S8. Re-examination of the EWAS data from the whole blood-derived DNA of NT1 patients analyzed with the 27K array. A.** Q-Q plot for sites with hypermethylation and hypomethylation in NT1 patients. **B.** Correlation between hypoSC and the degree of hypomethylation, adjusted for age. **C.** Correlation between the estimated values of GZMA and the degree of hypomethylation, adjusted for age. **D.** Correlation between hypoSC and the degree of hypomethylation, adjusted for age and the presence or absence of NT1. **E.** Correlation between the estimated values of GZMA and the degree of hypomethylation, adjusted for age and the presence or absence of NT1.

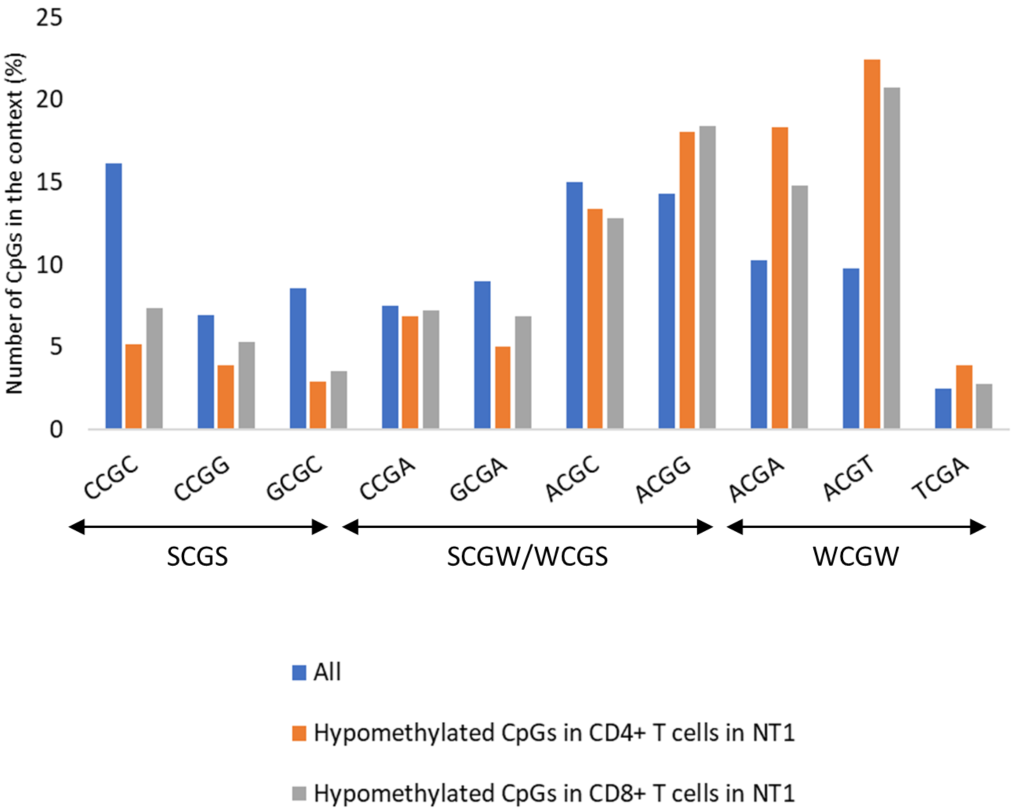


**Fig. S9. Examination of the sequences surrounding the analyzed CpG sites.** By classifying the types of base contexts adjacent to the analyzed CpGs, we compared NT1-related hypomethylated CpG sites to all analyzed CpG sites. W represents A and T, and S represents C and G.

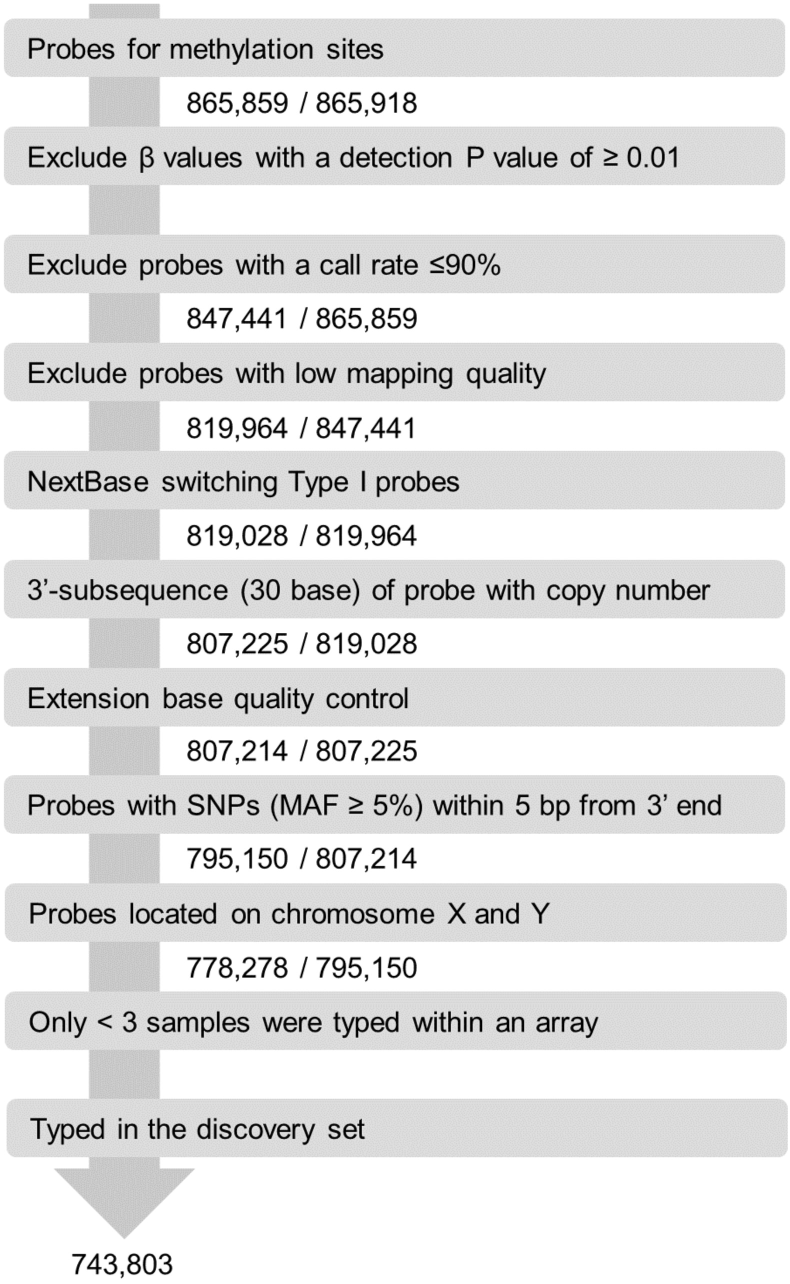


**Fig. S10. Procedure for filtering out the poor-performing probes in the replication set.** We used the same filtering procedure that was used for the discovery set, *i.e.,* we filtered out probes that have been reported to be poor-performing, those on sex chromosomes, and those that were typed in fewer than three samples in the array. We also focused on the probes that were typed in the discovery set.
